# *Gli2* and *Gli3* Regulate Horizontal Basal Cell-Mediated Regeneration of the Olfactory Epithelium

**DOI:** 10.1101/2022.08.29.504820

**Authors:** Anna Shirazyan, Haeyoung Park, Ariell M. Joiner, Justine Ra, Melissa S. Kim, Archana Kumari, Nicole E. Franks, Tien Peng, Robert K. Bradley, Charlotte M. Mistretta, Andrzej A. Dlugosz, Benjamin L. Allen

## Abstract

The olfactory epithelium (OE) is a specialized neuroepithelium that is replenished by two stem cell populations: globose basal cells (GBCs) and horizontal basal cells (HBCs). Previous work indicated that HBCs contain primary cilia, organelles that mediate Hedgehog (HH) pathway activity. However, a role for HH signaling in HBCs has not been investigated. We find that GLI2 and GLI3, transcriptional effectors of the HH pathway, are expressed in HBCs in the adult OE and that their expression expands following injury. Further, *Gli2*-expressing descendants contribute to all major OE cell types during OE regeneration. HBC-specific expression of constitutively active GLI2 drives inappropriate HBC proliferation, alters HBC identity, and culminates in a failure of HBCs to differentiate into olfactory sensory neurons (OSNs) following injury. HBC- specific deletion of endogenous *Gli2* and *Gli3* results in decreased HBCs and OSNs following OE injury. These data identify GLI2 and GLI3 as key regulators of HBC-mediated OE regeneration.

## Introduction

Olfactory dysfunction affects approximately 13.3 million older Americans (Hoffman et al., 2016). This can be due to factors that affect regeneration of the olfactory epithelium (OE), including physical injuries or pathogenic infections. The recent COVID-19 pandemic has resulted in loss of smell (anosmia) in patients (Aziz et al., 2021). To more effectively treat olfactory disorders, we need to better understand the signals that govern OE regeneration. The OE is a highly specialized neuroepithelium that contains olfactory sensory neurons (OSNs), which relay smell information (Graziadei and Graziadei, 1979). OSNs are vulnerable to the environmental toxins and pathogens present in the air, making the OE one of the few adult cites of neurogenesis (Graziadei and Graziadei, 1979; Morrison and Costanzo, 1989). Fortunately, two presumed stem cell populations can replenish OSNs: rapidly dividing globose basal cells (GBCs), and relatively quiescent horizontal basal cells (HBCs) (Schwob et al., 2017). HBCs lie in a monolayer along the OE basement membrane, with GBCs situated just above the HBCs. Both HBCs and GBCs can generate neuronal and non-neuronal progenitors, which give rise to immature OSNs (iOSNs), supporting glial-like Sustentacular cells (Sus), Bowman’s gland, and Microvillar cells (MVCs; (Schwob *et al*., 2017); Figure 1A). Although HBCs and GBCs mediate OE regeneration, the signals governing their function in homeostasis and injury-mediated repair are largely unexplored.

**Figure 1.**
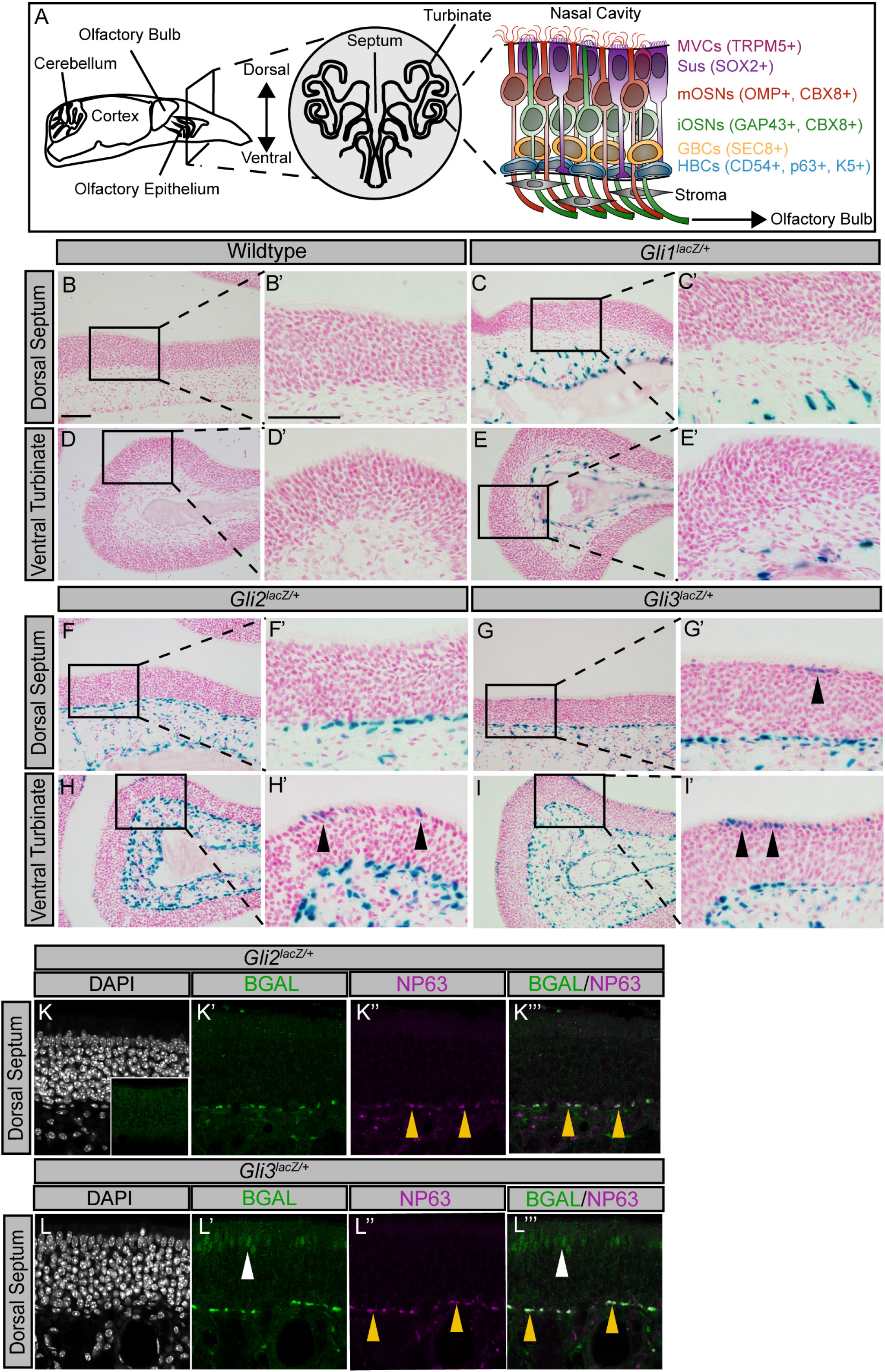
*Gli2* and *Gli3* are expressed in horizontal basal cells of the adult olfactory epithelium. (A) Cartoon of a sagittal view of the adult mouse head (left). Coronal view of a section through the adult mouse olfactory epithelium (middle). Schematic of OE cell types (right), including horizontal basal cells (HBCs; blue), globose basal cells (GBCs; orange), immature olfactory sensory neurons (iOSNs; green), mature olfactory sensory neurons (mOSNs; red), sustentacular cells (Sus; purple), and microvillar cells (MVCs; magenta). (B-I) X-gal staining of coronal sections from adult wildtype (B, B’, D, D’), *Gli1^lacZ/+^* (C, C’, E, E’), *Gli2^lacZ/+^* (F, F’, H, H’), and *Gli3^lacZ/+^* (G, G’, I, I’) reporter mice. Images are taken from both the dorsal septum (B-C’, F-G’) and a ventral turbinate (D-E’, H-I’). Black arrowheads indicate BGAL^+^ apical cells in *Gli2^lacZ/+^* (H’) and *Gli3^lacZ/+^* (G’. I’) mice. (J-M) Antibody detection of β-galactosidase (BGAL, green; J’, K’, L’, M’) and the HBC nuclear marker NP63 (magenta; J”, K”, L). DAPI denotes nuclei (gray; J, K, L, M). NP63 (magenta) and BGAL co-localization (green) in *Gli2^lacZ/+^* animals (K’’’; yellow arrowhead) and *Gli3^lacZ/+^* (L’’’; yellow arrowhead). Note that *Gli3*, but not *Gli2*, is expressed in apical cells at the septum of the adult OE (L’; L”’’; white arrowhead). Inset in K denotes the lack of BGAL detection in wildtype animals. Scale bars, B, B’, J = 50μm.

Hedgehog (HH) signaling is necessary for adult stem cell maintenance and function in many tissues, including multiple different epithelia (Petrova and Joyner, 2014). Notably, HH signaling is necessary for taste bud maintenance and renewal in the tongue, another chemosensory organ (Mistretta and Kumari, 2019). Targeted deletion of the HH transcription factor *Gli2* in the lingual epithelium results in taste bud loss and formation of atypical taste organs (Ermilov et al., 2016). Similar experiments with the HH pathway inhibitor LDE225 demonstrated a loss of taste buds and taste sensation in mice (Kumari et al., 2015; Kumari et al., 2017). Given the importance of HH signaling in lingual epithelial regeneration, it is possible that HH signaling is playing a similar role in another chemosensory epithelium – the olfactory epithelium. HH signaling is mediated by the primary cilium, a microtubule-based signaling center (Huangfu et al., 2003). In the absence of HH ligand, the canonical HH receptor Patched 1 (PTCH1) inhibits the activity of the GPCR-like protein Smoothened (SMO) resulting in phosphorylation and processing/degradation of GLI proteins, the transcriptional effectors of the HH pathway (Briscoe and Therond, 2013). In the presence of HH ligand, HH binding to PTCH1 results in de-repression of SMO, allowing SMO to accumulate in the cilium, resulting in GLIs accumulating at tips of primary cilia, and processing of GLIs into transcriptional activators that induce HH target gene expression (e.g. *Ptch1* and *Gli1*; (Briscoe and Therond, 2013)). GLI1 functions exclusively as a transcriptional activator and is also a target of HH signaling; GLI2 is typically the major transcriptional activator of the HH pathway; conversely GLI3 acts largely as a transcriptional repressor (Briscoe and Therond, 2013). GLI2 and GLI3 contain an N-terminal repressor domain and a C-terminal activator domain, and can be post-translationally processed into either their full length activator form or truncated repressor form (Sasaki et al., 1997; Sasaki et al., 1999). Though GLI2 and GLI3 are typically thought to have opposing roles, they can also play redundant roles depending on the context of the tissue (Chang et al., 2016; McDermott et al., 2005).

Previous work demonstrated that HBCs possess primary cilia and that HBC-specific ablation of primary cilia result in defective OE regeneration, indicating a functional role for primary cilia in HBCs (Joiner et al., 2015). Given that 1) HH signaling is a key regulator of tissue renewal across multiple epithelia, 2) HH signaling also is a key regulator of adult neurogenesis, and that 3) HH/GLI signaling rely on the primary cilium, a structure that is essential for HBC-mediated OE regeneration, we wondered whether HH signaling might play a role in stem cell-mediated adult OE regeneration.

Here, we show that the HH transcription factors *Gli2* and *Gli3* are expressed in HBCs and a subset of Sus cells in the OE. Additionally, we show that *Gli2+* HBCs can give rise to GBCs, OSNs, and Sus cells following injury. Further, we show that expression of constitutively active *Gli2* (GLI2A) in HBCs results in abnormal HBC proliferation and altered cell identity. Further, GLI2A-expressing HBCs fail to differentiate into OSNs following injury. Conditional *Gli2* and *Gli3* deletion in HBCs results in loss of HBCs and OSNs following injury. Together, these data suggest novel roles for GLI2 and GLI3 transcription factors in adult OE regeneration.

## Results

### Gli2 and Gli3 are expressed in HBCs and expand following OE injury

To investigate the expression of GLI transcription factors in the adult OE we utilized mice carrying *lacZ* reporter alleles for *Gli1^lacZ/+^* (Bai et al., 2002), *Gli2^lacZ/+^* (Bai and Joyner, 2001), and *Gli3^lacZ/+^* (Garcia et al., 2010). Specifically, we performed X-GAL staining on coronal sections, focusing on two different regions, the dorsal septum, and a ventral turbinate (Figure 1A). Like *Gli1*, *Gli2* is also expressed in the stroma underlying the OE; however, *Gli2* is more broadly expressed in both the proximal and distal stroma of the OE compared to *Gli1* (Figure 1F-F’). In contrast to *Gli1*, *Gli2* is also expressed in basal epithelial cells throughout the OE (Figure 1F-F’, H-H’). Further, *Gli2* is expressed in a subset of apical epithelial cells in the turbinates (Figure 1H-H’; black arrowheads). *Gli3* is also expressed broadly in the underlying stroma, and in basal cells throughout the OE (Figure 1G-G’, I-I’). Distinct from *Gli2*, *Gli3* is expressed in apical epithelial cells both at the septum (Figure 1G-G’; black arrowheads) and at turbinates (Figure 1I-I’;black arrowheads).

To define the cell types that expressed *Gli2* and *Gli3* in the OE, we used immunofluorescent antibody detection to mark *Gli*-expressing cells (BGAL^+^) and horizontal basal cells (HBCs; p63^+^) (Packard et al., 2011). While no BGAL signal was detected in WT animals (Figure 1K inset), in *Gli2^lacZ/+^* mice, BGAL co-localizes with NP63 (Figure 1K-K’’’; yellow arrow). Likewise, in *Gli3^lacZ/+^* mice, BGAL also co-localizes with p63 (Figure 1L-L’’’; yellow arrow), demonstrating that both *Gli2* and *Gli3* are expressed in HBCs of the adult OE. Co-staining with BGAL and a sustentacular (Sus) cell marker, SOX2 (Guo et al., 2010), revealed that BGAL co-localizes with apical SOX2^+^ cells in the ventral turbinates in both *Gli2^lacZ/+^* (Figure S1B-B’’’) and *Gli3^lacZ/+^* mice (Figure S1C-C’’’), indicating that *Gli2* and *Gli3* are expressed in distinct subsets of Sus cells in the OE. Together, these data demonstrate that *Gli2* and *Gli3*, but not *Gli1*, are expressed in the adult OE, displaying similar, but not identical expression patterns.

To further explore *Gli2* and *Gli3* expression in HBCs, a cell type that is quiescent during homeostasis (Fletcher et al., 2011; Packard *et al*., 2011), we sought to activate HBCs with an injury to the OE. Upon delivery of methimazole to the OE, Sus cells in the OE metabolize methimazole into a toxicant that results in the destruction of most of the OE (Genter et al., 1995). Notably, HBCs will be spared– they will proliferate and give rise to both neuronal and non-neuronal lineages, and at 8 weeks will fully regenerate the OE (Leung et al., 2007). We delivered 75mg/kg of methimazole to *Gli2^lacZ/+^* and *Gli3^lacZ/+^* via intraperitoneal (IP) injection, followed by analysis at 4 days of recovery post-injury a time point when HBCs are most active during regeneration (Packard *et al*., 2011). We performed *in situ* hybridization detection of *lacZ* transcripts, followed by immunofluorescent antibody detection of NP63 to mark HBC nuclei. Importantly, no *lacZ* expression is detected in wild-type uninjured and injured OE, demonstrating specificity of the *lacZ* probe (Figure 2A-C, D-F). Sparse *lacZ+* puncta are detected in NP63+ HBCs and stromal cells in uninjured *Gli2^lacZ/+^* (Figure 2G-I) and Gli3*^lacZ/+^* (Figure 2M-O) animals. At four days post-injury, however, *Gli2* expression is significantly increased throughout the entire OE (Figure 2J-L, S) as well as specifically in NP63^+^ HBCs (Figure 2U). Similarly, *Gli3* expression is also significantly increased in the OE following injury (Figure 2P-R, T) and in HBCs (Figure 2V). We confirmed these findings by performing X-GAL staining on *Gli2^lacZ/+^* and *Gli3^lacZ/+^* mice at 4 days and 8 weeks post methimazole injury (Figure 2S2 A-L’). Notably, X-GAL staining was not detected in WT animals at either timepoint (Figure S2 A-B’, G-H’). Similar to the *in situ* data, increased X-GAL staining was detected in *Gli2^lacZ/+^* mice at 4 days following injury, especially at ventral turbinates compared to the dorsal septum region (Figure S2C-D’). Since *Gli1* is a transcriptional readout for active HH signaling, we also used an endogenous *Gli1 in situ* probe to visualize *Gli1* expression prior to and following injury (Figure S1D-K). *Gli1* expression was not detected in the OE of uninjured mice, similar to X-GAL staining of *Gli1^lacZ/+^* animals (Figure S1E). No *Gli1* expression was detected in the OE 4 days following injury (Figure S1I), although stromal *Gli1* levels are significantly upregulated (Figure S1J-K). Taken together, these data suggest *Gli2* and *Gli3,* but not *Gli1,* are upregulated in the OE during early injury recovery.

**Figure 2.**
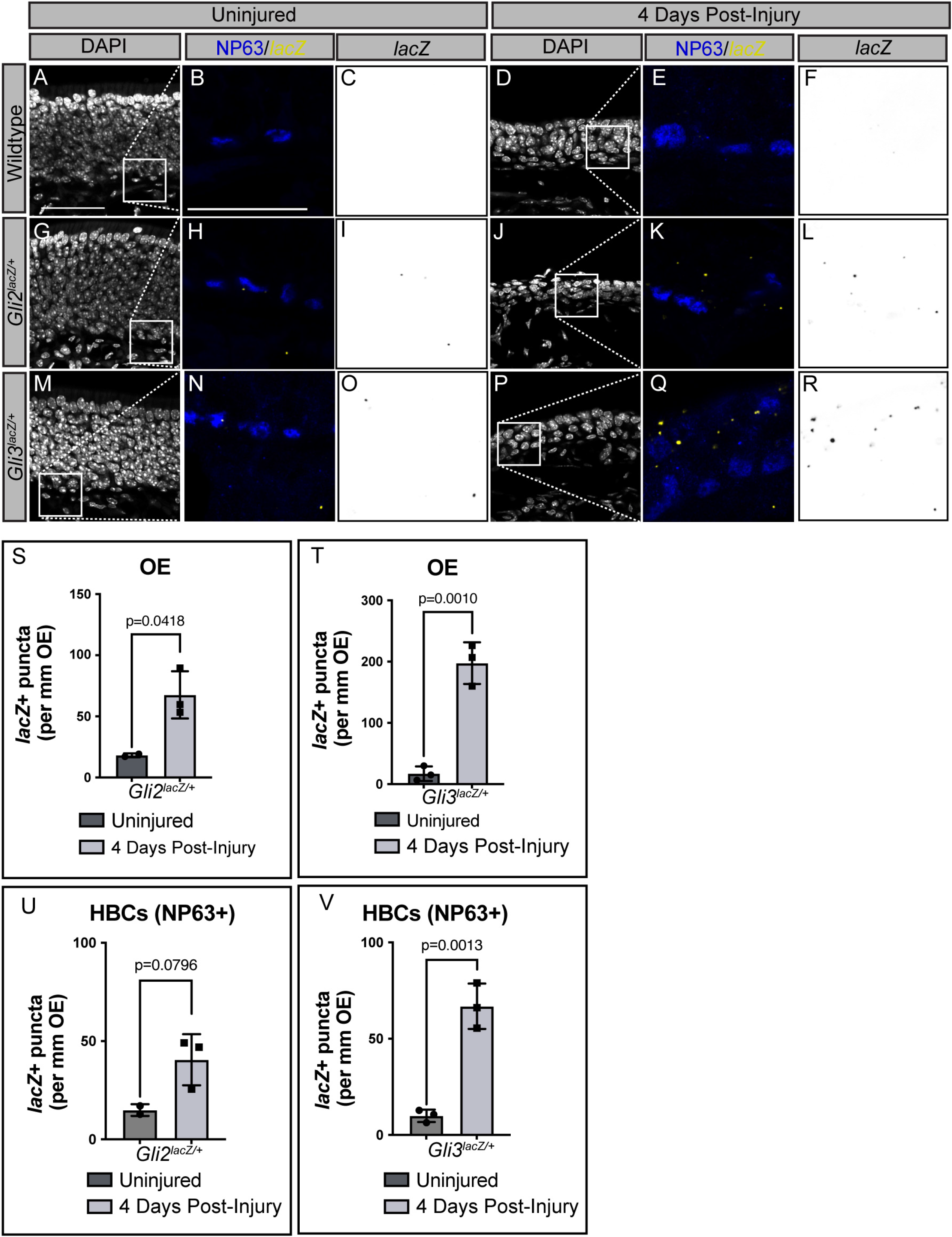
Selective upregulation of *Gli2* and *Gli3* expression in mice following methimazole-induced OE injury. (A-V) Adult WT, *Gli2^lacZ/+^* and *Gli3^lacZ/+^* reporter mice were either collected at 6-8 weeks of age or injured with a 75mg/kg IP injection of methimazole. Coronal sections were collected from the heads of either uninjured (A-C, G-I, M-O) or 4 days post methimazole injury mice (D-F, J-L, P-R). *In situ* hybridization detection of *lacZ* transcripts in sections from WT, *Gli2^lacZ/+^* and *Gli3^lacZ/+^* reporter mice prior to and following injury (yellow; B, H, N, E, K, Q). Inverted images of *lacZ* transcripts (black; C, I, O, F, L, R). Co-immunostaining with NP63 (blue; B, H, E, K, Q) demarcates HBCs. DAPI denotes nuclei (A, D, G, J, M, P), scale bar 50μm (A) and 25μm (B). Quantitation of *lacZ* puncta in the OE in *Gli2^lacZ/+^* (S) and *Gli3^lacZ/+^* (T) reporter mice prior and following methimazole injury. Quantitation of *lacZ+* puncta in HBCs from *Gli2^lacZ/+^* (U) and *Gli3^lacZ/+^* (V) reporter mice prior and following methimazole injury. N=2 uninjured *Gli2^lacZ/+^*, n=3 injured *Gli2^lacZ/+^*, n=3 uninjured *Gli3^lacZ/+^*, n=3 injured *Gli3^lacZ/+^*. Data are mean ± standard deviation. P-values were determined by a two-tailed unpaired t-test; n.s.= not significant (p>0.05). Scale Bar=50μm (A), 25μm (B).

### Gli2 descendants can give rise to all major OE cell types

Since *Gli2* expression in HBCs expands following injury, we employed a genetic lineage tracing approach to investigate the ability of *Gli2*-expressing cells and their descendants to contribute to different OE cell types. To achieve this, we bred *Gli2Cre^ER^* mice with a *ROSA26-lox-STOP-lox-tdtomato* reporter allele (Madisen et al., 2010; Wang et al., 2018). Coronal sections from the heads of mice treated with vehicle (corn oil) or tamoxifen (Figure 3A) were immunolabelled with antibodies directed against NP63 (HBCs) and SEC8 (GBCs). After either vehicle or tamoxifen treatment mice were rested for 72 hours then injected with methimazole to injure the OE (Figure 3A-G’’’’). We analyzed *Gli2*-expressing descendants at 8 weeks (full recovery) after injury by immunolabelling for different OE cell types- NP63/CD54 (HBCs), SEC8 (GBCs), SOX2 (Sus cells), and CBX8 (OSNs). Notably, no tdTomato+ progeny were visible in vehicle treated *Gli2Cre^ER^; tdTomato/+* mice (Figure 3B-B’’’’, D-D’’’’, F-F’’’’). We detected tdTomato+ labelled HBCs (Figure 3C’’, C’’’’; white arrowhead) and GBCs (Figure 3C’’’, C’’’’; yellow arrowhead) in fully recovered mice. Further, we detected tdTomato+ Sus cells (Figure 3E-E’’’’; blue arrowheads) and OSNs (Figure 3G- G’’’’; green arrowheads). These findings suggest that *Gli2+* HBCs can give rise to all major cell types of adult OE, namely GBCs, Sus cells, and OSNs.

**Figure 3.**
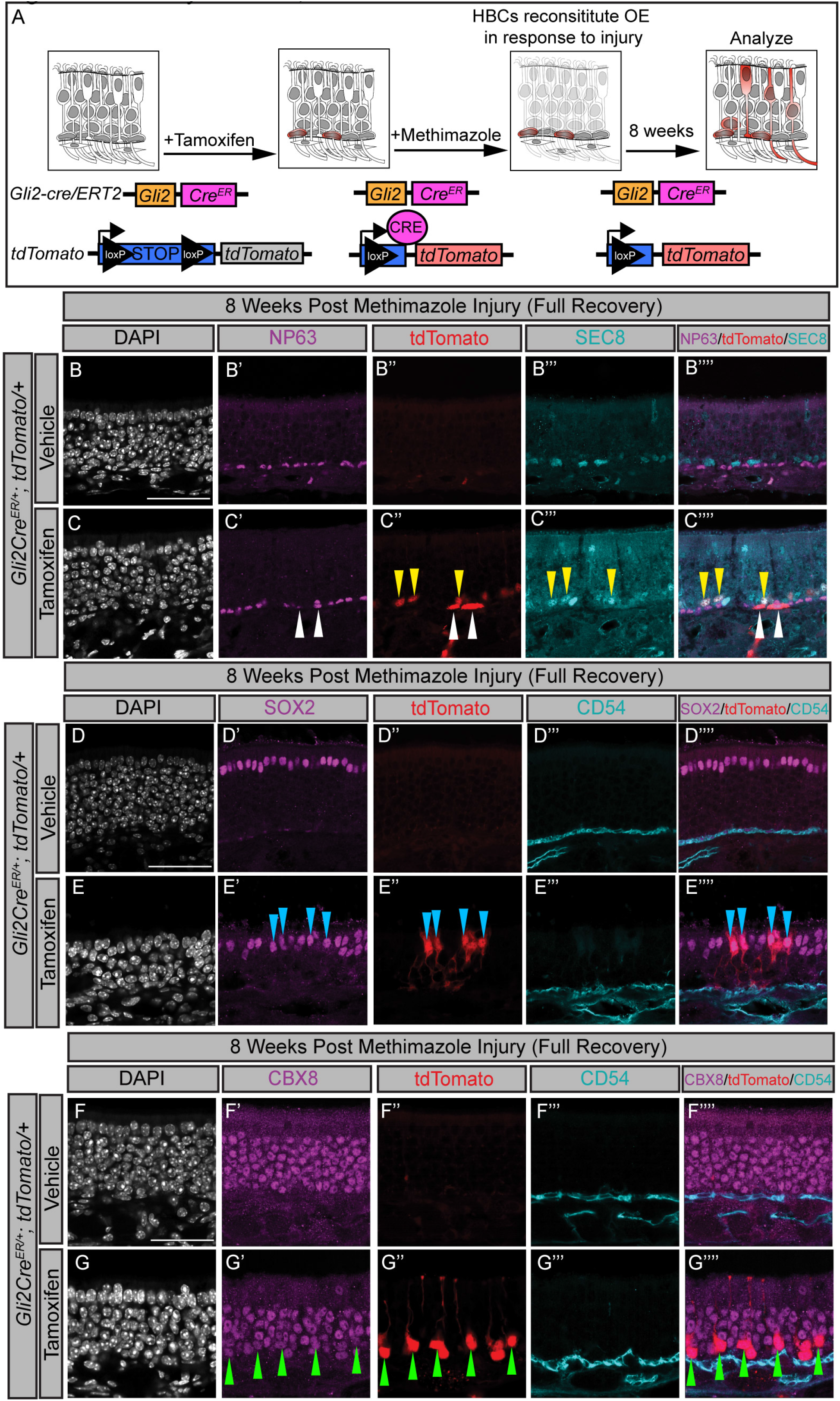
*Gli2*-descendants give rise to GBCs, Sus cells, and neurons following injury. (A) Cartoon of *Gli2* lineage tracing model using a tamoxifen inducible *Gli2Cre^ER^* mouse to mark *Gli2*-expressing cells. Mice carrying *Gli2Cre^ER^; tdTomato* alleles were given an IP injection with either vehicle (corn oil) or tamoxifen for 5 consecutive days at 100mg/kg. Upon tamoxifen administration, Cre recombinase excised the stop codon preceding the tdTomato gene, labelling *Gli2*-expressing cells and their descendants red. Mice were then injured with 75mg/kg methimazole and collected 8 weeks following injury (B-G’’’’). Coronal sections of vehicle- and tamoxifen-treated mice were stained with antibodies directed against NP63 label HBCs (magenta; B’, C’; white arrowheads) and SEC8 mark GBCs (cyan; B’’’, C’’’; yellow arrowheads). Antibodies directed against SOX2 label Sus cells apically (magenta; D’, E’; blue arrowheads) while antibodies directed against CD54 label HBCs (cyan; D’’’, E’’’, F’’’, G’’’). Antibodies directed against CBX8 denote neurons (magenta; F’, G’; green arrowheads). Cells originating from a *Gli2* lineage are labelled with tdTomato (red; B’’, C’’. D’’, E’’, F’’, G’’). Merged images of NP63, tdTomato and SEC8 (B’’’’, C’’’’), SOX2, tdTomato, CD54 (D’’’’, E’’’’), and CBX8, tdTomato, and CD54 (F’’’’, G’’’’). DAPI denotes nuclei (gray; B, C, D, E, F, G), scale bar (B, D, F) = 50μm.

### GLI2 activator drives HBC proliferation and alters HBC identity

Given that increased *Gli2* expression correlates with HBC activation and that *Gli2* descendants contribute to all major OE cell types, we wondered whether the primarily transcriptional activator function ascribed to GLI2 in other systems (Bai *et al*., 2002) might drive HBC activation in the OE. To test this notion, we utilized a doxycycline inducible system to express a constitutively active form of GLI2 lacking the N-terminal repressor domain (GLI2ΔN) in HBCs. We crossed mice that contain a reverse tetracycline transactivator driven by a Keratin 5 promoter (Krt5rtTA) (Diamond et al., 2000) to mice containing a tet operon inducible MYC-tagged *GLI2ΔN* transgene (Grachtchouk et al., 2011) (Figure 4). Mice were fed a doxycycline diet to induce *GLI2ΔN* expression in HBCs and analyzed at 1 day, 3 days, 5 days, and 8 days following induction (Figure 4B-G’’’’, Figure S3A-D). Due to the aggressive nature of driving overactive HH signaling these mice do not survive past 7-8 days (Grachtchouk *et al*., 2011). Coronal sections of heads from control and experimental mice were stained with antibodies directed against MYC (GLI2ΔN), CD54 (HBCs), and Ki67 (actively cycling cells). While no MYC expression was detected in vehicle-treated *Krt5rtTA;GLI2ΔN* mice (Figure 4B’), MYC immunostaining was detected in HBCs of *Krt5rtTA;GLI2ΔN* mice treated with doxycycline for 1 day (Figure 4C’). Immunostaining for Ki67 (Figure 4B’’, C’’), a marker for actively cycling cells, identifies GBCs lying apical to CD54+ HBCs (Figure 4B’’’, C’’’; white arrowheads) but not HBCs themselves, indicating that at 1 day post doxycycline induction HBCs are not actively proliferating. In contrast, at 3 days post doxycycline induction MYC^+^ HBCs (Figure 4E’, E’’’; yellow arrowheads) express Ki67 (Figure 4E’’; yellow arrowheads), while control HBCs do not (Figure 4D-D’’’’). At 5 days post doxycycline induction MYC^+^ HBCs (Figure 4G’, G’’’; yellow arrowheads) continue expressing Ki67 (Figure 4G’’; yellow arrowheads), while control HBCs do not (Figure G-G’’’’). Notably, GLI2ΔN protein is detected in HBC-associated primary cilia, an important organelle for HH signal transduction, at 8 days following doxycycline administration (Figure S4 F-J; white arrowhead). HBCs reach peak proliferation at 3 days post doxycycline induction (Figure S3 C-D), resulting in significantly increased HBC number at 3, 5, and 8 days post doxycycline induction (Figure S3 A, D). These data suggest that GLI activator can induce HBC proliferation.

**Figure 4.**
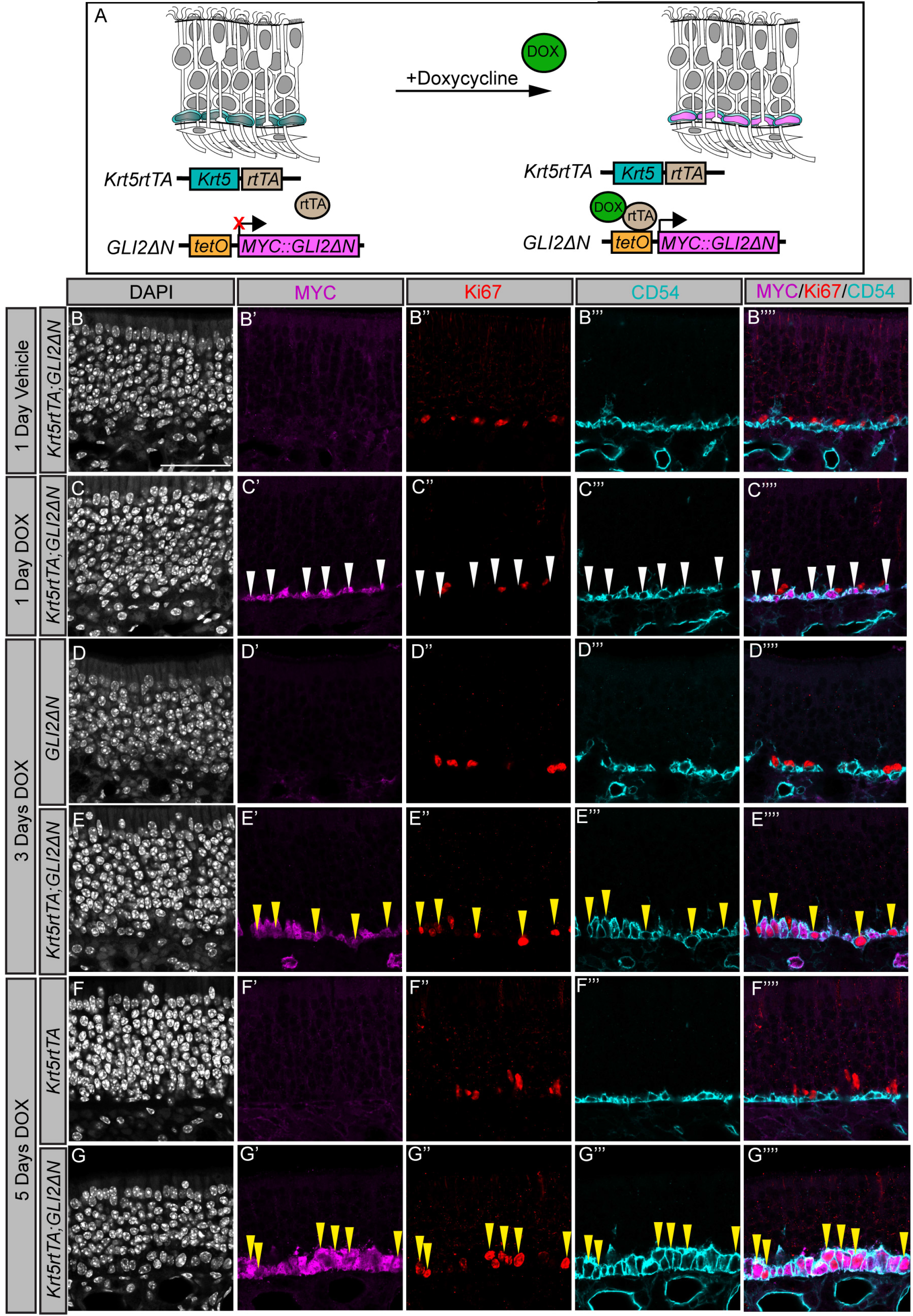
Constitutively active GLI2 drives HBC proliferation. (A) Cartoon of *GLI2ΔN* transgene activation using a doxycycline-inducible mouse model. Upon doxycycline (DOX, green) administration, mice carrying a Keratin 5 reverse tetracycline transactivator (*Krt5rtTA*, blue) transgene and a tet-regulated MYC-tagged *GLI2ΔN* transgene (pink) express constitutively active *GLI2ΔN* specifically in HBCs of the OE by the rtTA (brown) protein binding at the tet operon (oragnge). (B-G’’’’) Antibody detection of MYC (magenta, B’-G’), Ki67 (red, B’’-G’’), and CD54 (cyan, B’’’-G’’’) at 1 Day Vehicle treatment (B-B’’’’), 1 Day DOX treatment (C-C’’’’), 3 days DOX treatment (D-D’’’’, E-E’’’’), and 5 days DOX treatment (F-F’’’’, G-G’’’’). DAPI denotes nuclei (gray; B-G). Merged MYC, Ki67, and CD54 images (B’’’’-G’’’’). MYC::GLI2ΔN protein is detectable at 1 day following DOX treatment (magenta, C’, C’’’’; white arrowheads). Following 3 days of DOX administration HBCs express Ki67 (red, E’’, E’’’’; yellow arrowheads). Ki67 expression in HBCs persists following 5 days of DOX administration (red, G’’, G’’’’; yellow arrowheads). Scale bar (B) = 50μm.

To further characterize the consequences of HBC-specific GLI2A expression on the OE, we examined markers for other cell types, starting with GBCs. Strikingly, and in contrast to control mice (Figure 5A-E), at 8 days post doxycycline induction HBCs co-express SEC8, a pan GBC marker (Joiner *et al*., 2015) (Figure 5 F-M). The total number of HBCs (Figure 5K) is significantly increased while the total number of GBCs remains unchanged (Figure 5 L). Further, we performed immunostaining for SOX2, which labels a subset of HBCs and GBCs basally and Sus cells apically in control mice (Figure 5N-Q). GLI2A-expressing HBCs also co-express SOX2 (Figure 5R-V). Immunolabelling for SOX9 (normally a marker for Bowman’s gland cells) also revealed extensive co-labelling with HBCs in *Krt5rtTA;GLI2ΔN* mice (Figure S4O-R), compared to control mice (Figure S4K-N, S). Interestingly, *Krt5rtTA;GLI2ΔN* HBCs do not co-express the neuronal marker CBX8 (Figure S4T), and there was no significant change in neurons in *Krt5rtTA;GLI2ΔN* OE (Figure S4U). In contrast, there is a subtle but significant increase in Sus cell number in mice expressing GLI2A (Figure 5W). Taken together, these data suggest that constitutive GLI activator function drives altered HBC identity and increased Sus cells.

**Figure 5.**
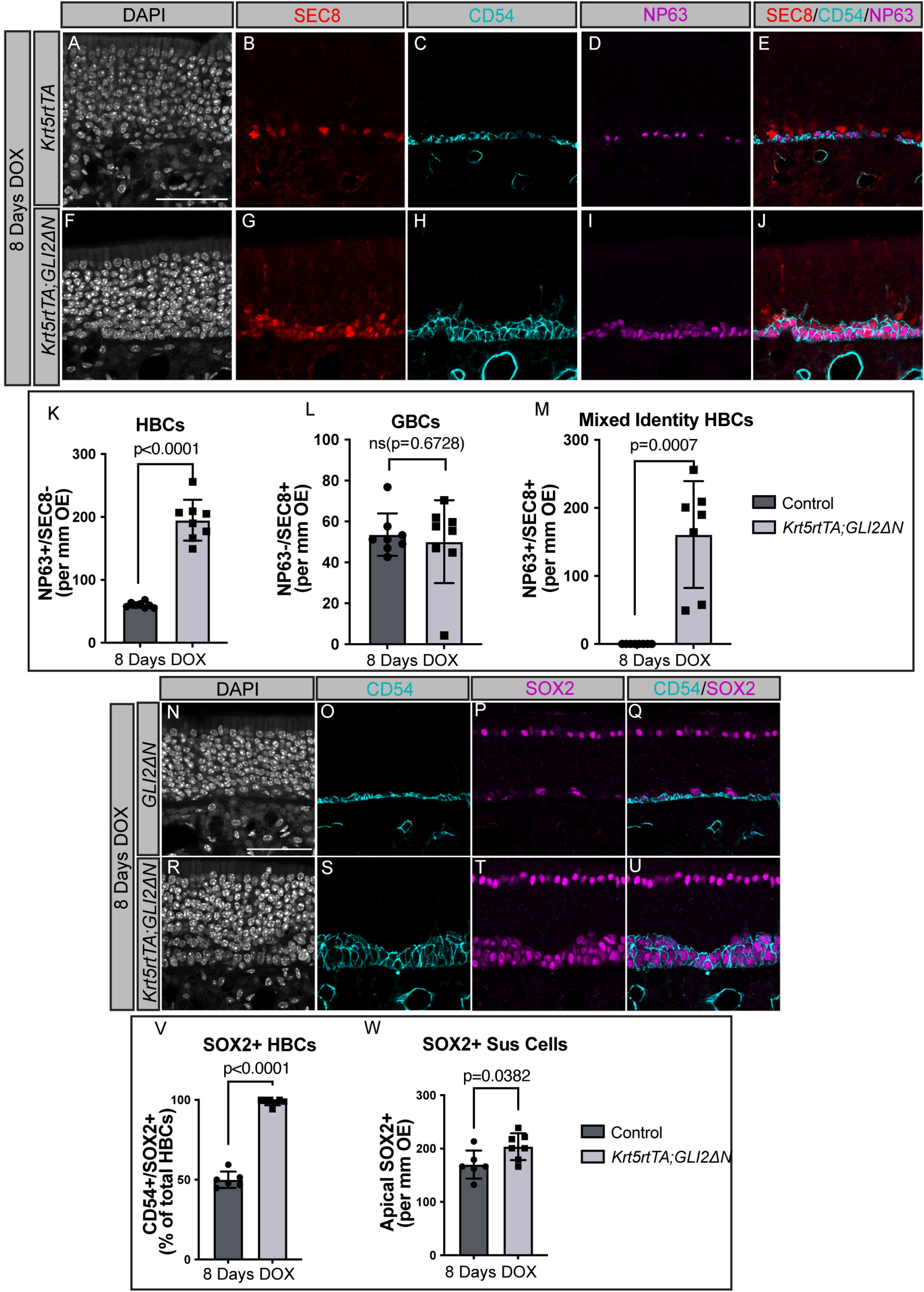
Constitutively active GLI2 alters HBC identity. (A-J) Coronal sections from adult mice treated with DOX for 8 days were analyzed with immunostaining using different OE markers. Antibodies directed against SEC8 (red; B, G) identify GBCs, while antibody detection of CD54 (cyan; C,H) and NP63 (magenta; D,I) denotes HBCs. DAPI indicates nuclei (gray; A, F). Merged SEC8, CD54, and NP63 images (E, J). Quantitation of HBCs (NP63^+^, SEC8^-^; K), GBCs (NP63^-^,SEC8^+^; L), and mixed identity HBCs (NP63^+^, SEC8^+^; M). n=8 control and n=8 *Krt5rtTA*;*GLI2ΔN* animals analyzed (K-M). (N-U) Coronal sections from adult mice treated with DOX for 8 days were analyzed with immunostaining using different OE markers. Antibody detection of CD54 (cyan; O,S) and apical SOX2 (Sus cell marker, magenta; P,T). DAPI indicates nuclei (N, R). Merged CD54 and SOX2 images (Q,U). Quantitation of SOX2^+^ HBCs (SOX2^+^, CD54^+^; V) and SOX2^+^ Sus cells (Apical SOX2^+^ cells; W). n=6 control and n=7 *Krt5rtTA*; *GLI2ΔN* animals analyzed (V-W). Scale bars (A, N) = 50μm. Control refers to mice containing either the *Krt5rtTA* or *GLI2ΔN* transgene. Data are mean ± standard deviation. P-values were determined by a two-tailed unpaired t-test; n.s.= not significant (p>0.05).

### GLI2 activator promotes HBC differentiation to Sus Cell, but not OSN lineages following OE injury

Next, we assessed the effect of GLI2A on HBC function. Since HBCs are primarily quiescent in homeostatic conditions (Fletcher *et al*., 2011; Packard *et al*., 2011), we sought to activate them by injuring the OE. After inducing control and *Krt5rtTA;GLI2ΔN* mice with doxycycline for 24 hours, we injured the OE with an IP injection of methimazole at 75mg/kg (Figure 6, Figure S5). Mice were maintained on a doxycycline diet following injury and allowed to recover for 7 days (Figure 6A). We first analyzed the basal cells in control and GLIA mice following methimazole injury (Figure S5A-N). Coronal sections of mice collected at 7 days following injury were stained with antibodies for different OE markers in addition to Ki67 to mark actively cycling cells. While only a small fraction of HBCs in control OE are actively cycling (less than 5%) proliferative (Figure S5A-E, K), GLI2A-expressing HBCs are significantly more proliferative (Figure S5F-K). As a consequence there is a significant increase in HBC number in injured *Krt5rtTA;GLI2ΔN* mice (Figure S5L). In contrast, GBC number is significantly downregulated (Figure S5M). As with uninjured animals, we also observed an upregulation in SOX9+ cells in injured GLI2A-expressing animals (Figure S5N).

**Figure 6.**
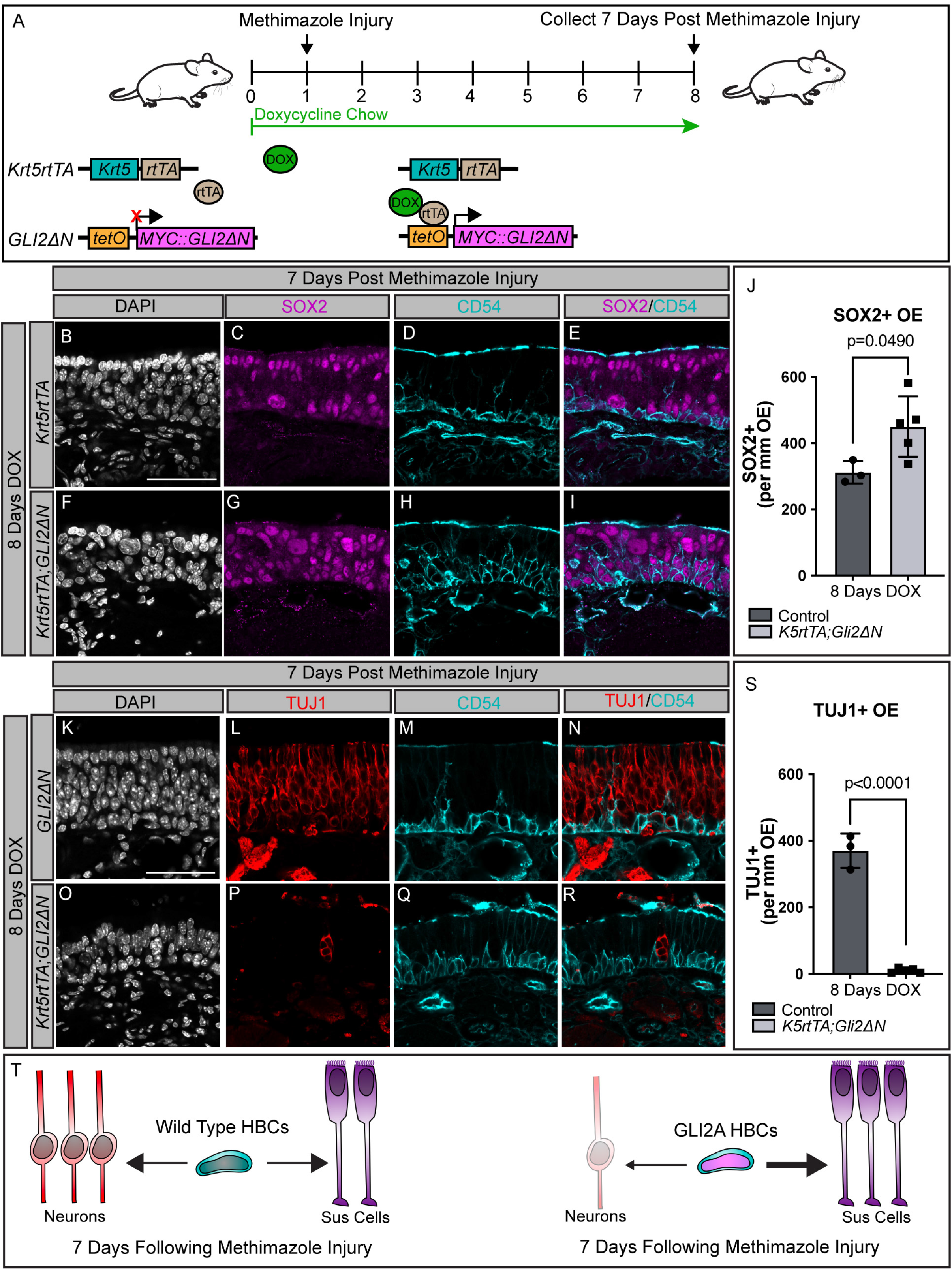
GLI2A-expressing HBCs fail to contribute to neuronal lineages following injury. (A) Cartoon of *GLI2ΔN* transgene activation using a doxycycline-inducible mouse model. Upon doxycycline (DOX, green) administration, mice carrying a Keratin 5 reverse tetracycline transactivator (*Krt5-rtTA*, blue) transgene and a tet-regulated MYC-tagged *GLI2ΔN* transgene (pink) express constitutively active *GLI2ΔN* specifically in HBCs of the OE. 1 day following doxycycline induction, mice are injured with a 75mg/kg intraperitoneal (IP) injection of methimazole. Following methimazole injury, mice are kept on doxycycline and collected 7 days following injury. (B-S) Coronal sections from adult mice treated with DOX for 8 days, 7 days following methimazole injury, were analyzed with immunostaining using different OE markers. Antibodies directed against SOX2 (magenta; C, G) denote apical lying Sus cells and a subset of basal HBCs and GBCs, while antibody detection of CD54 (cyan; D, H, M, Q) exclusively identifies HBCs. Merged SOX2 and CD54 images (E, I). DAPI indicates nuclei (gray; B, F, K, O). Quantitation of the percentage of SOX2^+^ cells in the OE (J). Antibodies directed against TUJ1 (red; L, P) denote neurons. Merged TUJ1 and CD54 images (N, R). Quantitation of the percentage of TUJ1^+^ cells in the OE (S). Scale bar (B, K), 50μm. n=3 control and n=5 *Krt5rtTA*; *GLI2ΔN* animals analyzed. Control refers to mice containing either the *Krt5rtTA* or *GLI2ΔN* transgene. Data are mean ± standard deviation. P-values were determined by a two-tailed unpaired t-test; n.s.= not significant (p>0.05). (T) Summary of HBC-mediated OE reconstitution at 7 days following injury in wild type (left) and *Krt5rtTA*;*GLI2ΔN* mice (right).

To investigate the consequences of HBC-specific GLI2A expression on OE differentiation, including the formation of neurons and Sus cells, we first immunostained coronal sections of control and *Krt5rtTA;GLI2ΔN* mice with SOX2 to mark Sus cell progenitors. We discovered that GLI2A-expressing HBCs give rise to a significantly higher amount of total SOX2^+^ progenitors, in comparison to control HBCs (Figure 6J). The OE in control HBCs also appeared more organized with a clear layer of early apical Sus cells separate from basal cells (Figure 6B-E), whereas the OE in *Krt5rtTA;GLI2ΔN* mice appeared much more disorganized with expanded SOX2 staining (Figure 6F-I). Additionally, we immunolabelled for neurons using TUJ1 (Figure 6K-S).

Surprisingly, there was a significant decrease in TUJ1^+^ progenitors in GLI2A OE compared to control OE (Figure 6S). GLI2A-expressing HBCs give rise to very few neurons (Figure 6O-R) compared to control HBCs which generate a well-defined TUJ1^+^ layer of cells (Figure 6K-N). In short, while wild-type HBCs can give rise to both neurons and Sus cells following injury, HBCs with overactive HH signaling primarily give rise to Sus cells and not neurons (Figure 6T). This suggests that GLI activator not only drives HBC proliferation, but also drives HBCs preferentially toward a Sus cell lineage.

### GLI2 and GLI3 are necessary for proper OE regeneration following injury

To investigate the function of endogenous GLIs in HBCs we utilized a conditional deletion mouse model to selectively delete *Gli2* and *Gli3* in HBCs. We crossed in an HBC-specific, tamoxifen-inducible *Krt5- Cre^ER^* (Diamond *et al*., 2000) to *Gli2^fl/fl^* (Corrales et al., 2006) and *Gli3^fl/fl^* (Blaess et al., 2008) alleles surrounded by loxP sites. A *ROSA26-lox-STOP-LOX-tdTomato* (Madisen *et al*., 2010) allele was also crossed in to track HBC lineages following methimazole injury. Mice were treated with 100mg/kg of tamoxifen for 5 consecutive days, thus generating *Gli2^CKO^*, *Gli3^CKO^*, and *GLI2/3^CKO^* mice lacking either *Gli2* or *Gli3* individually, or combination in HBCs. We then performed methimazole injury at 75mg/kg to stimulate HBC proliferation and differentiation. We collected and analyzed HBC progeny in *Gli2^CKO^* (Figure S6A-E), *Gli3^CKO^* (Figure S6F-J), and *GLI2/3^CKO^* (Figure 7) mice 8 weeks following methimazole injury (full recovery). We used the tdTomato reporter to track HBC-derived cells and performed immunolabeling for different OE markers to determine the effect of HBC-specific conditional *Gli* deletion on OE recovery. *Gli2^CKO^* OE are comparable to control OE across all cell types. We observed no significant difference in the numbers of HBCs (Figure S6B), GBCs (Figure S6C), OSNs (Figure S6D), or Sus cells (Figure S6E). In contrast, *Gli3^CKO^* animals display subtle but significant increases in HBCs (Figure S6G), GBCs (Figure S6H), and Sus cells (Figure S6J). The number of OSNs are slightly but not significantly decreased (Figure S6I). Given that GLI3 largely functions as a transcriptional repressor in other tissues (Petrova et al., 2013; Sasaki *et al*., 1997; Sasaki *et al*., 1999; Wang et al., 2000), the increase in basal and Sus cells in *Gli3^CKO^* OE complement our findings with HBC-specific constitutive GLI2 activator expression (cf. Figure 6).

**Figure 7.**
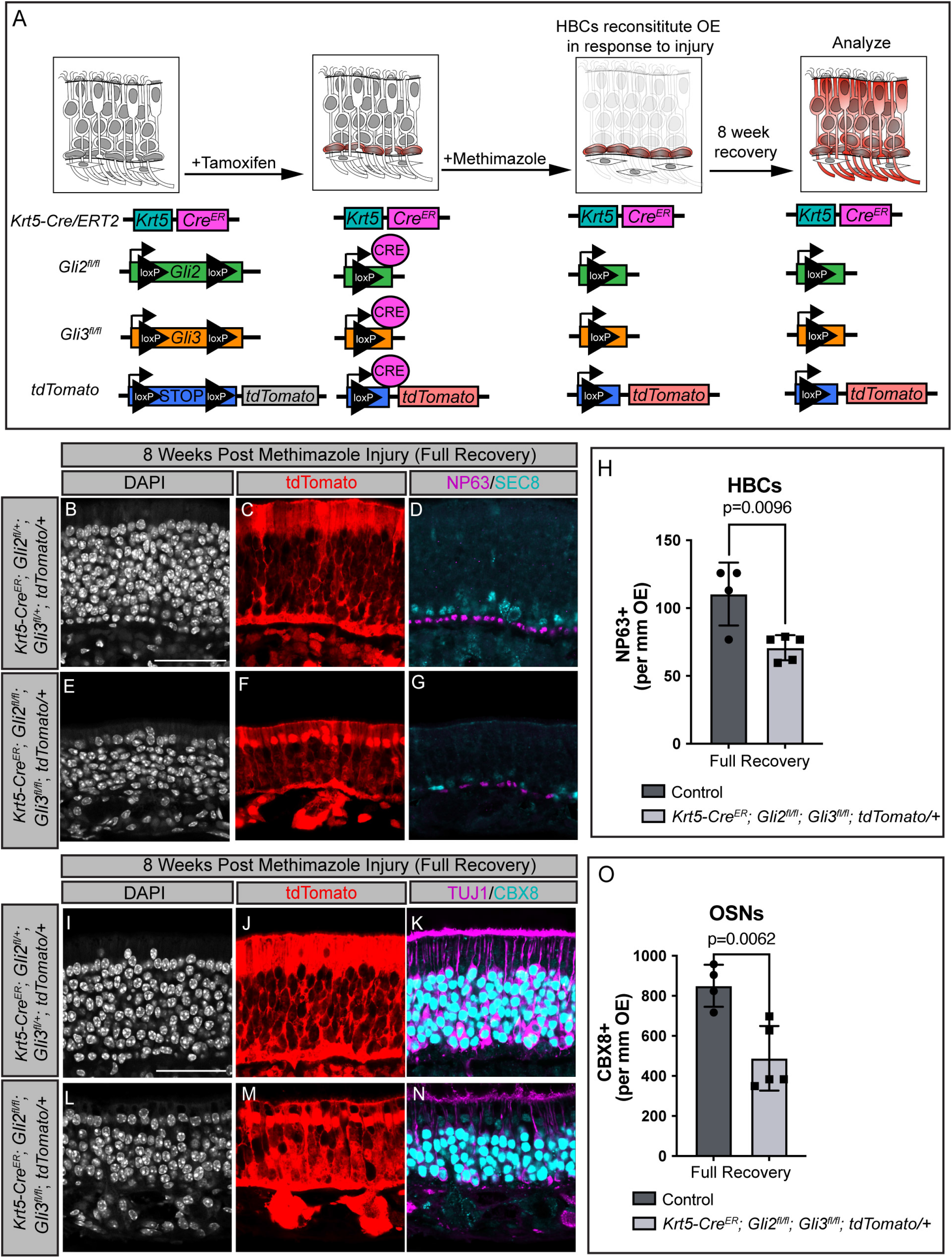
Simultaneous HBC-specific *Gli2* and *Gli3* deletion results in defective OE regeneration. (A) Cartoon of tamoxifen-inducible HBC-specific *Gli2* and *Gli3* deletion. Mice carrying a *Krt5-Cre^ER^* allele were crossed with mice containing *Gli2^fl/fl^* and *Gli3^fl/fl^* alleles as well as a *tdTomato* reporter allele. Tamoxifen administration induces Keratin 5 (*Krt5*)-expressing cells to drive *Gli2* and *Gli3* deletion and *tdTomato* expression. After tamoxifen administration (I.P., 100mg/kg for 5 consecutive days), mice were rested for 72 hours, then injured with methimazole (I.P., 75mg/kg) and allowed to recover for 8 weeks. (B-O) Coronal sections from adult mice collected 8 weeks following injury were analyzed with immunostaining for various OE markers. Antibodies directed against NP63 (magenta; D, G) mark HBCs, while antibodies directed against SEC8 (cyan; D, G) denote GBCs. Antibodies directed against Tuj1 (magenta; K, N) and CBX8 (cyan; K, N) mark OSNs. DAPI denotes nuclei (gray; B, E, I, L), scale bar (B, I), 50μm. Quantitation of HBC (H) and OSN (O) number per mm of OE. n=4 control and n=5 *Krt5-Cre^ER^*; *tdTomato/+*; *Gli2^fl/fl^*; *Gli3^fl/fl^* animals analyzed. Endogenous tdTomato reporter immunofluorescence (red; C, F, J, M) demarcates HBCs and their lineages. Control animals refers to *Krt5-Cre^ER^*;*tdTomato/+*;*Gli2^fl/+^*;*Gli3^fl/+^* mice or mice lacking the *Krt5-Cre^ER^* allele (H, O). Data are mean ± standard deviation. P-values were determined by a two-tailed unpaired t-test; n.s.= not significant (p>0.05).

Since *Gli2* and *Gli3* can have redundant functions (McDermott *et al*., 2005), we sought to delete *Gli2* and *Gli3* simultaneously in HBCs (*GLI2/3^CKO^*) (Figure 7). We first confirmed efficient deletion of *Gli2* and *Gli3* in our genetic mouse model using *in situ* hybridization prior to methimazole injury (Figure S7A-F). *Gli2* expression was 86% downregulated in *GLI2/3^CKO^* OE (Figure S7C, E) compared to control mice (Figure S7A, E). *Gli3* was 74% downregulated in *GLI2/3^CKO^* OE (Figure S7D, F) compared to control mice (Figure S7B, F). *GLI2/3^CKO^* animals display a range of OE phenotypes following injury- while some regions are grossly disturbed and thin (Figure S7G, I) others looked normal (Figure S7H, J). Quantitation of basal cell numbers in *GLI2/3^CKO^* OE revealed that HBCs are significantly decreased (Figure 7E-H), compared to control OE (Figure 7B-D, H). GBC numbers are also subtly decreased, although not significantly (Figure 7E-G, Figure S7R). Next, we co-immunolabelled for TUJ1 and CBX8 to mark OSNs (Figure 7I-O). We discovered that OSNs are significantly decreased in *GLI2/3^CKO^* OE (Figure 7L-N, O), compared to control OE (Figure 7I-K, O). Notably, we observed no change in Sus cell number (Figure S7K-Q), even in more significantly perturbed areas (Figure S7N inset). These findings suggest that combined HBC-specific *Gli2* and *Gli3* deletion results in reduced HBC numbers and a subsequent failure to properly regenerate the OE following injury, particularly with regard to OSN differentiation. Taken together, these data illustrate novel individual and combined roles for GLI transcription factors in HBC-mediated OE regeneration by regulating the proliferation and differentiation of HBCs following injury (Figure 7S).

## Discussion

Here, we explored the contribution of the HH pathway to OE regeneration. Specifically, we found that *Gli2* and *Gli3*, encoding two key transcriptional effectors of the HH pathway, are expressed in HBCs and distinct subset of Sus cells in the OE. Further, we discovered that *Gli2* and *Gli3* expression expands following OE injury. Constitutive GLI2A expression in HBCs results in HBC hyperproliferation and defective differentiation following OE injury. These data suggest that transient GLI activation following OE injury allows for HBC activation and proliferation, but that abrogation of GLI activator function is necessary for subsequent HBC differentiation. HBC-specific conditional *Gli2* deletion does not result in any detectable phenotype, while HBC-specific conditional *Gli3* deletion results in increased HBC, GBC and Sus cell numbers. Simultaneous conditional *Gli2* and *Gli3* deletion in HBCs results in improper OE regeneration and significantly decreased HBC, GBC, and OSN numbers. Together, these data illustrate novel roles for GLI signaling in HBC function and demonstrate that GLI2 and GLI3 are necessary for OE regeneration post-injury.

### Diversity of HH response across different types of epithelia

HH signaling plays important but diverse roles in many adult regenerative epithelia. Two such examples are the lingual and respiratory epithelium. HH signaling is essential for proper maintenance and function of taste organs in the lingual epithelium (Mistretta and Kumari, 2019). SHH ligand is present in the stem cells and nerves that innervate the taste bud (Ermilov *et al*., 2016; Liu et al., 2013). As in the OE, *Gli2* is also expressed in basal cells of the lingual epithelium as well as in the underlying stroma (Liu *et al*., 2013). *Gli1* is also present in the lingual stroma and epithelium, which differs from the OE where it is expressed exclusively in the stroma (Ermilov *et al*., 2016). GLI2A expression in Keratin5+, *Gli2*+ cells in the tongue results in aberrant proliferation of lingual epithelial cells (Liu *et al*., 2013). These findings are remarkably like the OE, where GLI2A-expressing HBCs become hyperproliferative. Further, the lingual epithelium is dependent on active HH signaling for maintenance. Chemical HH pathway inhibition using a SMO antagonist LDE225 results in disrupted taste organs and taste sensation (Kumari *et al*., 2015). Further studies in the OE with chemical HH pathway blockade can elucidate if canonical HH signaling is necessary for OE maintenance and regeneration. HH signaling is also a key component of adult lung regeneration and maintenance (Wang et al., 2019). During lung homeostasis, SHH ligand in the epithelium signals to surrounding GLI1+ mesenchymal cells (Peng et al., 2015). Genetic ablation of *Shh* results in mesenchymal cell proliferation and an increase in epithelial cells in the airway (Peng *et al*., 2015). Additionally, genetic activation of HH signaling in the lung mesenchyme (through expression of oncogenic *Smo*) results in an emphysema-like phenotype (Wang *et al*., 2018). Thus, HH signaling appears to have a restrictive role and maintains quiescence in the lung. Conversely, HH signaling is necessary for HBC proliferation and can drive proliferation in the olfactory epithelium. The adult lingual, respiratory, and olfactory epithelia are all regenerative, diverse, and unique structures. The differential deployment of HH/GLI signaling in each of these epithelial appears to be one mechanism by which a single pathway can contribute to the maintenance and regeneration of related, but distinct tissues.

### Regulation of GLI expression and function in the olfactory epithelium

Our data indicate an increase in *Gli2* and *Gli3* expression following injury in the OE. In contrast, there is no detectable expression of the HH target gene *Gli1* in the OE with *Gli1* expression remains solely stromal even following OE injury. These data suggest potentially non-canonical roles for GLI2 and GLI3 in the OE. While a definitive source of HH-producing cells in the OE remains to be elucidated, one potential source includes SHH in nerves that innervate the OE (Kumari et al., 2017). SHH ligand is also present in the olfactory bulb (OB), particularly in the glomeruli (Gong et al., 2009). Further, OSNs are able to respond to SHH ligand *in vitro* (Gong et al., 2009). Since OSNs directly connect the OE with the OB, it is possible that SHH is delivered to HBCs and the underlying stroma through OSN axonal projections. Another promising source of ligand is IHH, which is expressed in bone and regulates proliferation and differentiation of chondrocytes (Vortkamp et al., 1996). The OE and underlying stroma are directly in contact with the bone in the turbinates, making the turbinates a potential source of IHH. Further careful investigation into *Shh* and *Ihh* expression in the OE and surrounding tissues is necessary to elucidate if canonical HH signaling is regulates *Gli2* and *Gli3* in the OE.

Since HH target gene expression is not detected in the OE, it is possible that GLI2 and GLI3 function independently of HH ligands. A possible avenue of GLI regulation in the OE is through the NP63 transcription factor. NP63 is an important regulator of stemness in HBCs as illustrated in (Packard *et al*., 2011). NP63 maintains HBC quiescence and transiently turns off following MeBr gas injury to the OE (Packard *et al*., 2011). Our data illustrate an upregulation of *Gli2* and *Gli3* immediately following injury to the OE. It is possible that NP63 could negatively regulate *Gli2* and *Gli3* during OE homeostasis, resulting in upregulation of *Gli2* and *Gli3* when NP63 levels decrease following injury. Previous studies in mammary cancer stem cells illustrated that NP63 can directly regulate HH pathway genes such as SHH and GLI2 by binding to upstream regulatory regions (Memmi et al., 2015). Additionally, our data indicates that driving GLI2A results in significant upregulation of NP63. This suggests a possible feedback loop between GLIs and NP63 expression. Careful investigation of whether NP63 can bind at *Gli2* and *Gli3* loci in HBCs can further our understanding of GLI regulation in HBCs.

### Crosstalk between Hedgehog and Notch signaling

While the primary focus of this study is on HH signaling and GLI transcription factors, other signaling pathways have been described in the adult OE. Specifically, Notch signaling has been implicated in OE regeneration (Schwob *et al*., 2017). Importantly, Notch signaling pathway components are present in HBCs, along with Sus cells and underlying stroma (Herrick et al., 2018; Herrick et al., 2017). Previously work demonstrated that modulation of various Notch signaling pathway components altered HBC cell fate following OE injury (Herrick *et al*., 2018). These findings echo our observations in the OE following GLI2A overexpression in HBCs. Similarly, in our HH overactivation model HBC cell fate is altered to Sus/non-neuronal following injury. It is possible that HH and Notch signaling coordinate to dictate HBC cell fates in response to injury. Along these lines, previous studies have demonstrated that Notch and HH signaling can work in conjunction in the context of development. For example, Notch signaling directs HH morphogen activity in the developing ventral spinal cord (Kong et al., 2015). Additionally, the HH co-receptor GAS1 directly binds to NOTCH1 in the developing forebrain, and coordinates HH and Notch signaling in the neuroepithelium (Marczenke et al., 2021). Future studies examining Notch signaling pathway components in our genetic models can elucidate cooperation between HH and Notch signaling in HBCs.

### Limitations of Study

This study used multiple gain-of-function and loss-of-function mouse genetic approaches to investigate the roles of GLI2 and GLI3 in HBCs during OE regeneration. Nevertheless, we could broaden our study by including human tissue in our analysis. Previous studies have successfully isolated single cell transcripts from mOSNs of human patients (Durante et al., 2020). A similar approach could be utilized by sorting HBCs and examining HH pathway transcripts in human patients. Understanding the role of HH signaling in human OE can also open therapeutic avenues for patients with loss of smell.

Though our work focused primarily on HBCs and regeneration, we also discovered that *Gli2* and *Gli3* are expressed in other cell types in the OE as well. For example, GLI2 and GLI3 are regionally expressed in Sus cells in the OE, opening a potential avenue of study of HH signaling in Sus cells. There is a precedent for key developmental pathways playing a role in Sus cell maintenance – for example genetic ablation of Notch2 results in Sus cell death in post-natal OE (Rodriguez et al., 2008). It is possible that genetic ablation of *Gli2* and *Gli3* specifically in Sus cells could affect Sus cell function. Additionally, it is possible that Sus cells and HBCs directly communicate through HH signaling. Similar studies have shown expression of both Notch ligand and receptors in HBCs and Sus cells, suggesting possible cell-cell communication (Herrick *et al*., 2017). Using a similar approach, we could study the expression of additional HH pathway components in Sus cells to assess the potential for juxtracrine signaling between HBCs and Sus cells. Overall, a closer study of HH signaling in Sus cells would broaden our understanding of Sus cell function in the adult OE.

The functional role of the underlying stroma, also known as the lamina propria, during OE injury remains largely unexplored. The stroma is a highly vascularized tissue where fibroblasts, dendritic bundles from OSNs, and blood vessels are all present. During infection, the stroma is an important source of inflammatory cytokines and defense against invading parasites (Imamura and Hasegawa-Ishii, 2016). Studies have examined the stroma in chronic inflammatory rhinosinusitis models of mice and demonstrated that the immune system can directly affect HBCs and the OE (Chen et al., 2019). Thus, the immune response could be playing a role in OE regeneration following methimazole-induced injury. Notably, recent work from our lab indicate that GLIs regulate the immune response in the context of pancreatic cancer (Scales et al., 2022). In this study we determined that all three *Glis* are expressed in the stroma underlying the OE. These data raise the question of GLI-dependent stromal contributions to maintenance and regeneration of the OE.

## Acknowledgements

We thank members of the Allen lab for their feedback and technical support throughout this study. We thank Dr. Bradley Goldstein (Duke University) for technical advice and valuable feedback on the OE injury experiments. We acknowledge the Department of Cell and Developmental Biology at the University of Michigan and the O’Shea laboratory for access to research equipment. We acknowledge the Biomedical Research Core Facilities Microscopy Core for access to and maintenance the microscopes used for this study. This work was supported by a Ruth L. Kirschstein Predoctoral Individual National Research Service Award (F31 DC018206 to A.S.), Bradley M. Patten Fellowship (to A.S.), Rackham Merit Fellowship (to A.S.), and the Hearing, Balance, and Chemical Senses Training Program (T32 DC00011, University of Michigan to A.S.).

This work was also supported by the National Institutes of Health (R01 DC014428 to B.L.A., R.K.B., A.A.D., C.M.M.).

## Author Contributions

Conceptualization, A.S., A.M.J., and B.L.A.; Methodology, A.S., A.M.J., and N.E.F., and B.L.A.; Investigation, A.S., H.P., A.M.J., J.R., M.S.K., and A.K.; Formal Analysis, A.S.; Writing – Original Draft, A.S.; Writing – Review & Editing, A.S. and B.L.A.; Funding Acquisition, A.S., R. B., A. A. D., C.M.M., and B.L.A.; Resources, T.P., A.A.D., and B.L.A.; Supervision, B.L.A.

## Materials and Methods

### Key Resources Table

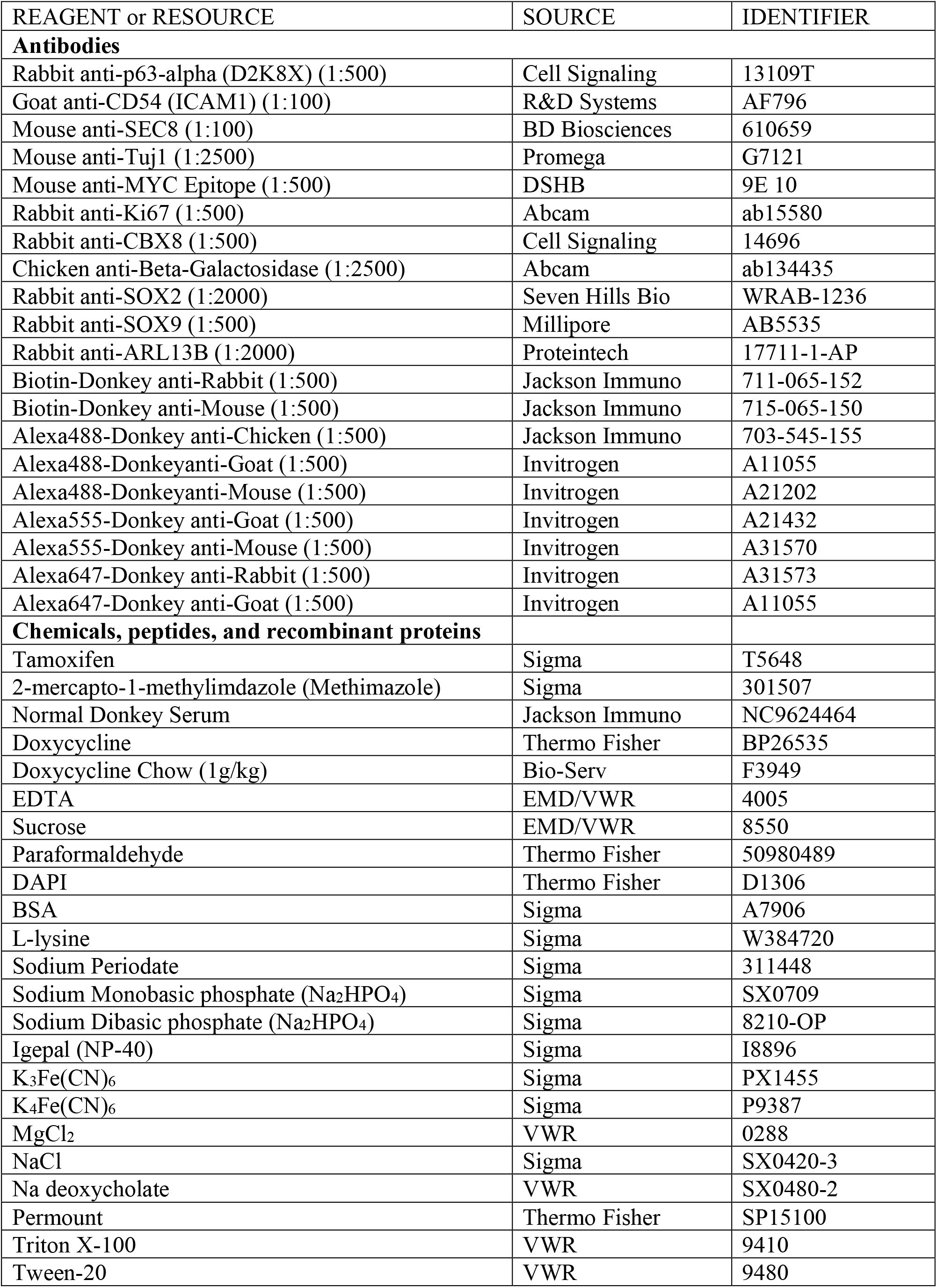
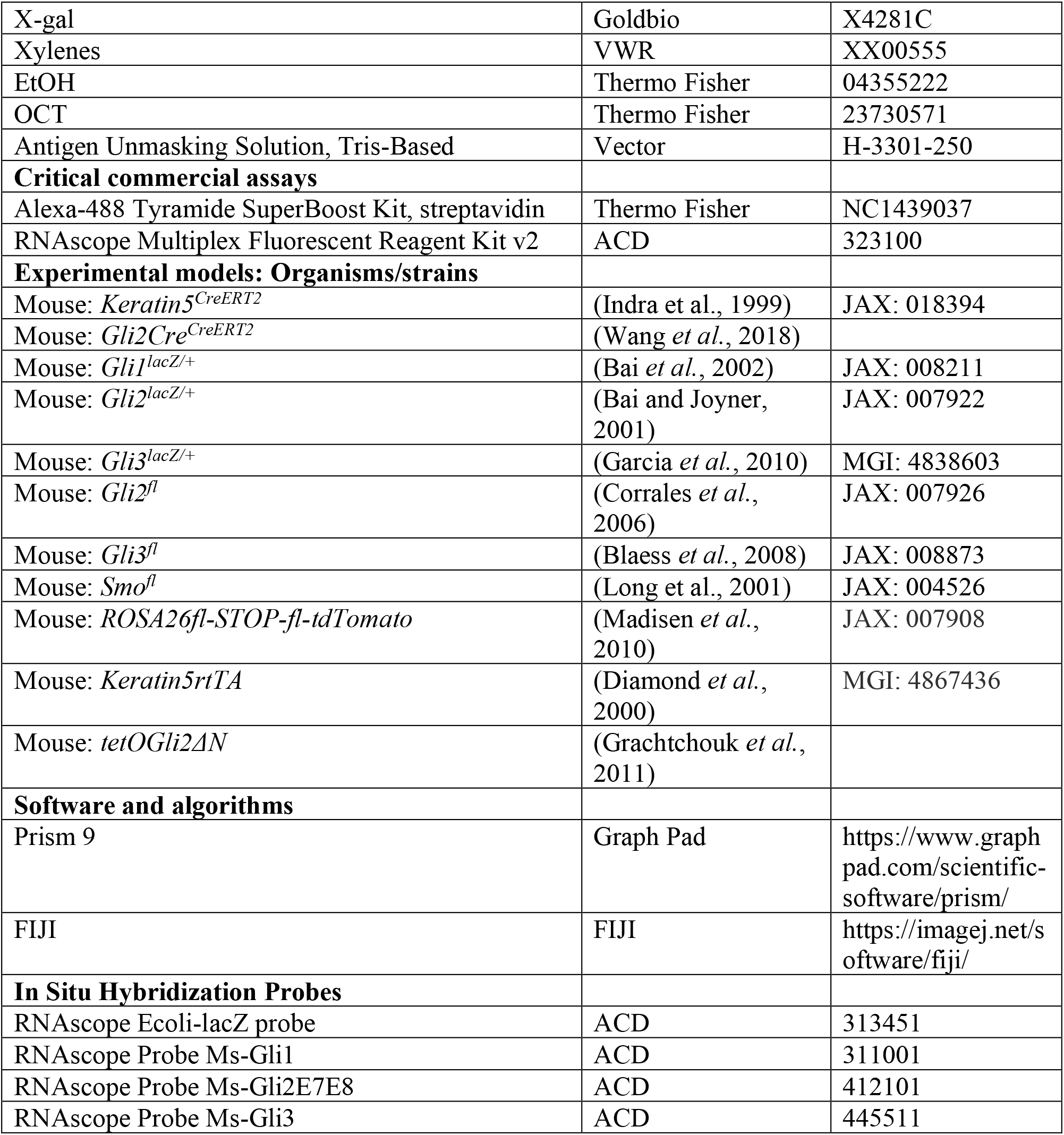

### Animals and breeding

All mice were maintained on a mixed BL/6, 129, and CD1 genetic background. *Gli1^lacZ/+^*{Bai, 2002 #3}, *Gli2^lacZ/+^* (Bai and Joyner, 2001), and *Gli3^lacZ/+^* (Garcia *et al*., 2010) mice have all been described previously. *Keratin5rtTA;tetOGli2ΔN* (Grachtchouk et al., 2000) mice were provided by Dr. Andrzej Dlugosz (University of Michigan, Ann Arbor MI). *Gli2Cre^ERT2^* mice were provided by Dr. Tien Peng (University of California, San Francisco). HBC-specific *Gli2* and *Gli3* deletion was accomplished by breeding animals carrying a *Keratin5Cre^ERT2^* allele with mice carrying *Gli2* and *Gli3* alleles flanked by loxP sites. To lineage trace progeny from HBCs that underwent *Cre*-mediated recombination, we crossed in a *ROSA26fl-STOP-fl-tdTomato* allele (JAX: 007908) into all conditional deletion mouse lines (*Gli2^fl^* and *Gli3^fl^*). For all experiments male and female adult mice (6-8 weeks of age) were used. All animal procedures were reviewed and approved by the Institutional Animal Care and Use Committee (IACUC) at the University of Michigan.

### Tamoxifen preparation and administration

Tamoxifen was dissolved in sterile corn oil at 55°C for approximately 2-3 h at 40mg/mL with occasional agitation until in solution. For all experiments with conditional deletion mice (*ROSA26fl-STOP-fl-tdTomato*, *Gli2^fl^*, and *Gli3^fl^*) mice were injected 100mg/kg of tamoxifen intraperitoneally (I.P.) for 5 consecutive days. Mice were allowed to rest for 72 h prior to methimazole lesion (described below).

### Doxycycline preparation and administration

Doxycycline powder was dissolved at 200μg/ml in 5% sucrose in autoclaved water and administered to mice through drinking water. *Keratin5rtTA;tetOGli2ΔN* mice were given doxycycline water during first three days of doxycycline chow treatment. Doxycycline chow (1g/kg, Bio-Serv #F3949) was administered to *Keratin5rtTA;tetOGli2ΔN* mice at 6-8 weeks until date of euthanasia.

### Methimazole lesion

Methimazole (2-mercapto-1-methylimdazole) was dissolved in sterile 1X PBS and administered to control and experimental mice through an intraperitoneal (IP) injection at 75mg/kg following either tamoxifen or doxycycline treatment.

### X-gal staining

Mice were anesthetized with 30% isoflurane, transcardially perfused with 2% PLP solution (2% paraformaldehyde, 0.01M sodium periodate, 0.01M monobasic and dibasic phosphates, and 90mM L-lysine as described in (Packard *et al*., 2011), and decapitated. Heads were post-fixed in 2% PLP for 1hr at 4 °C. Tissue was then decalcified in 0.5 M EDTA overnight at 4°C; cryoprotected in 10% (1h), 20% (1h), and 30% sucrose overnight at 4°C; and frozen in OCT compound. Coronal sections of the olfactory epithelium and olfactory bulb (OB) were cut at 12μm thickness on a Epredia™ Microm HM525 NX Cryostat. Sections were stored at -80 °C until the day of staining. Beta-galactosidase activity was detected with X-gal staining solution (5mM K3Fe(CN)6, 5mM K4Fe(CN)6, 2mM MgCl2, 0.01% Na deoxycholate, 0.02% NP-40, 1mg/mL X-gal). Sections were incubated with X-gal staining solution overnight in 37 °C. After staining, sections were washed 3 x 5 min with 1x PBS, pH 7.4, counterstained with nuclear fast red for 5 min and dehydrated in an ethanol series (70% ethanol, 95% ethanol, 100% ethanol and 100% Xylenes) followed by application of coverslips with permount mounting media. Sections were visualized on a Nikon E-800 Upright Widefield Microscope.

### Immunohistochemistry

Mice were anesthetized with 30% Fluriso (isoflurane, VetOne), transcardially perfused with 4% paraformaldehyde (PFA), and decapitated. Heads were post-fixed in 4% PFA for 20–24 h at 4°C. Tissue was then decalcified in 0.5 M EDTA overnight at 4°C; cryoprotected in 10% (1h), 20% (1h), and 30% sucrose overnight at 4°C; and frozen in OCT compound. 12ìm thick coronal sections of the olfactory epithelium and olfactory bulb (OB) were collected on a Epredia™ Microm HM525 NX Cryostat. Sections were stored at -80°C until day of staining. On the day of staining, sections were baked at 70°C for 10 min, washed 3 x 5 min with 1X PBST (0.01% Triton X), pH 7.4, then immediately placed in TRIS antigen retrieval solution for 15 minutes in a 92°C water bath. Sections were then blocked with 10% donkey serum for 1h at RT then incubated with primary antibody overnight at 4°C. The next day sections were washed with 3 x 5 min with 1X PBST (0.01% Triton X), pH 7.4, then incubated with secondary antibody for 1h at RT, For some antibodies (SEC8, CBX8) TSA amplification was used to detect signal. Nuclei were labeled with DAPI for 10 min at room temperature (RT) and slides were mounted with coverslips using Immu-mount aqueous mounting medium. Sections were visualized on a Leica upright SP5X confocal microscope.

### *In situ* hybridization

Mice were anesthetized with 30% Fluriso (isoflurane, VetOne), transcardially perfused with 10% neutral buffered formalin (10% NBF), and decapitated. Heads were post-fixed in 10% NBF for 24 h at RT. Tissue was then decalcified in 0.5 M EDTA overnight at 4°C; cryoprotected in 10% (1h), 20% (1h), and 30% sucrose overnight at 4°C; and frozen in OCT compound. Coronal sections of the olfactory epithelium and olfactory bulb (OB) were cut at 12μm thickness on a Epredia™ Microm HM525 NX Cryostat. Sections were stored at -80 °C until the day of staining when sections were baked at 70°C for 10 min then washed 5 min in 1X PBST (0.01% Triton X), pH 7.4. Fluorescent *in situ* hybridization was performed using ACD RNAscope Multiplex Fluorescent Reagent Kit v2, according to manufacturer’s instructions (ACD 323100-USM). Pretreatment conditions were optimized for the olfactory epithelium: antigen retrieval was performed for 15 min, and protease treatment was performed for 1 min. Following the RNAscope assay, slides were incubated in primary antibody and followed the immunohistochemistry staining protocol described above. Sections were visualized on a Leica upright SP5X confocal microscope.

### Quantitation and Statistical analysis

All the data are represented as mean ± standard deviation. All statistical analyses were performed using GraphPad statistic calculator (GraphPad Software, La Jolla California USA, www.graphpad.com). Statistical significance was determined using two-tailed Student’s t-test. Significance was defined according GraphPad Prism style: non-significant (*p*>0.05) and significant (*p*≤0.05). For all the experimental analyses a minimum of 3 mice of each genotype were examined; each n represents a mouse. To quantify cell counts all images were blinded and four separate areas of the OE (dorsal septum, dorsal turbinate, ventral septum, and ventral turbinate) were averaged per n. All the statistical details (statistical tests used, statistical significance and exact value of each n) for each experiment are specified in the figure legends.

## Supplemental Figure Legends

**Figure S1.**
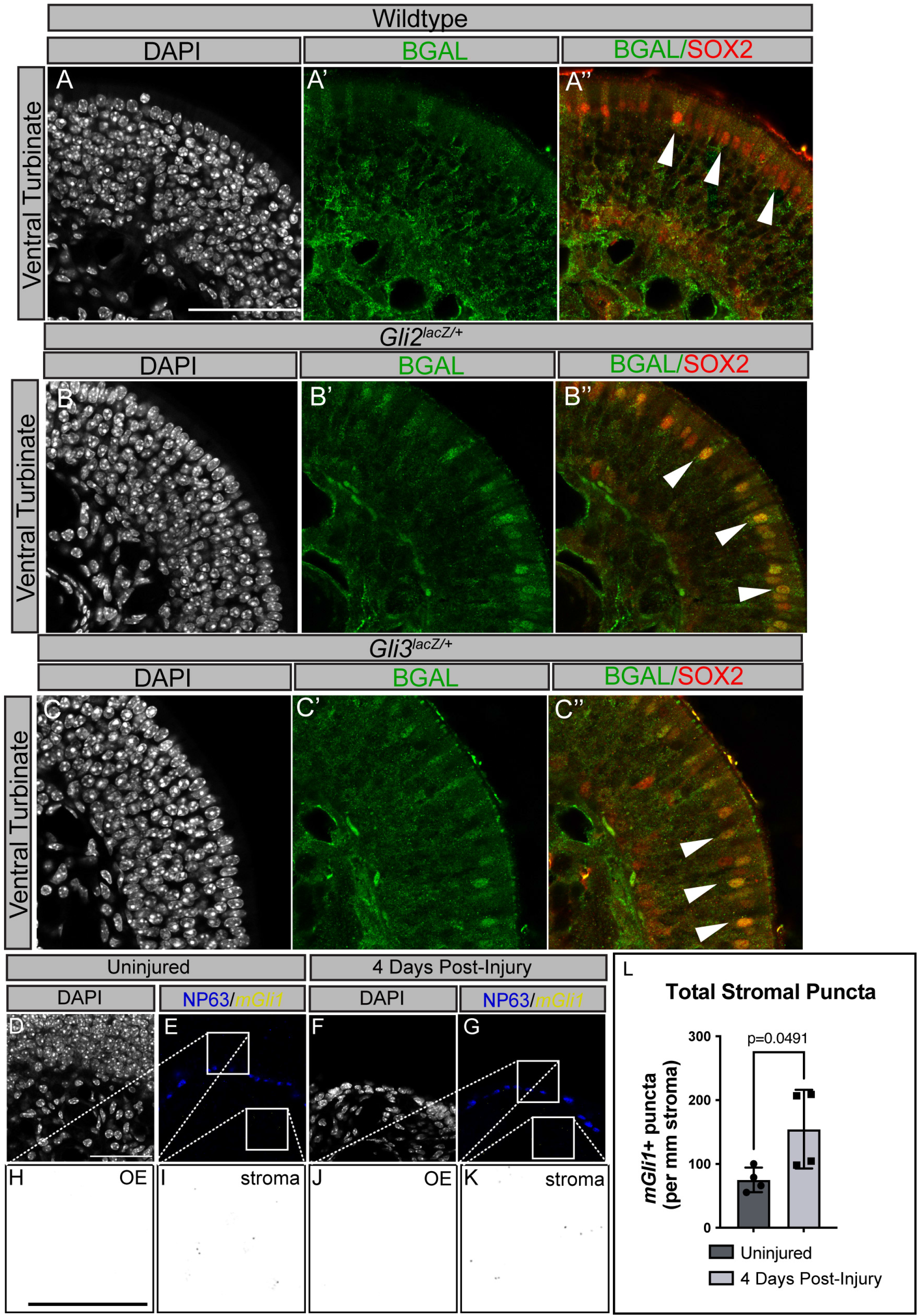
*Gli2* and *Gli3* are expressed in turbinate-associated sustentacular cells, while *Gli1* expression is stromally restricted, even following OE injury. **(A-B)** Coronal sections of adult WT, *Gli2^lacZ/+^*, and *Gli3^lacZ/+^* mice were stained with antibodies against β-galactosidase (green; A’-C’) and a marker for Sus cells, SOX2 (red; A’’-C’’). DAPI denotes nuclei (gray; **A-C**). Scale bar= 50μm. (**D-K**) Adult mice were either collected at 6-8 weeks of age or injured with a 75mg/kg IP injection of methimazole. Coronal sections of either uninjured OE or OE 4 days following injury were processed for *in situ* hybridization with a probe to detect endogenous mouse *Gli1* (yellow; **E, G**). Antibodies directed against NP63 demarcate HBCs (blue; E, G). Inverted images of the *mGli1* probe in OE (black; H, J) and stroma (black; I, J). DAPI denotes nuclei (**D, F**), scale bars, 50μm (A, D), 25μm (H). Quantitation of *mGli1* puncta in the stroma underlying the OE, n=4 uninjured, n=4 injured (K). Data are mean ± standard deviation. P-values were determined by a two-tailed unpaired t-test; n.s.= not significant (p>0.05).

**Figure S2.**
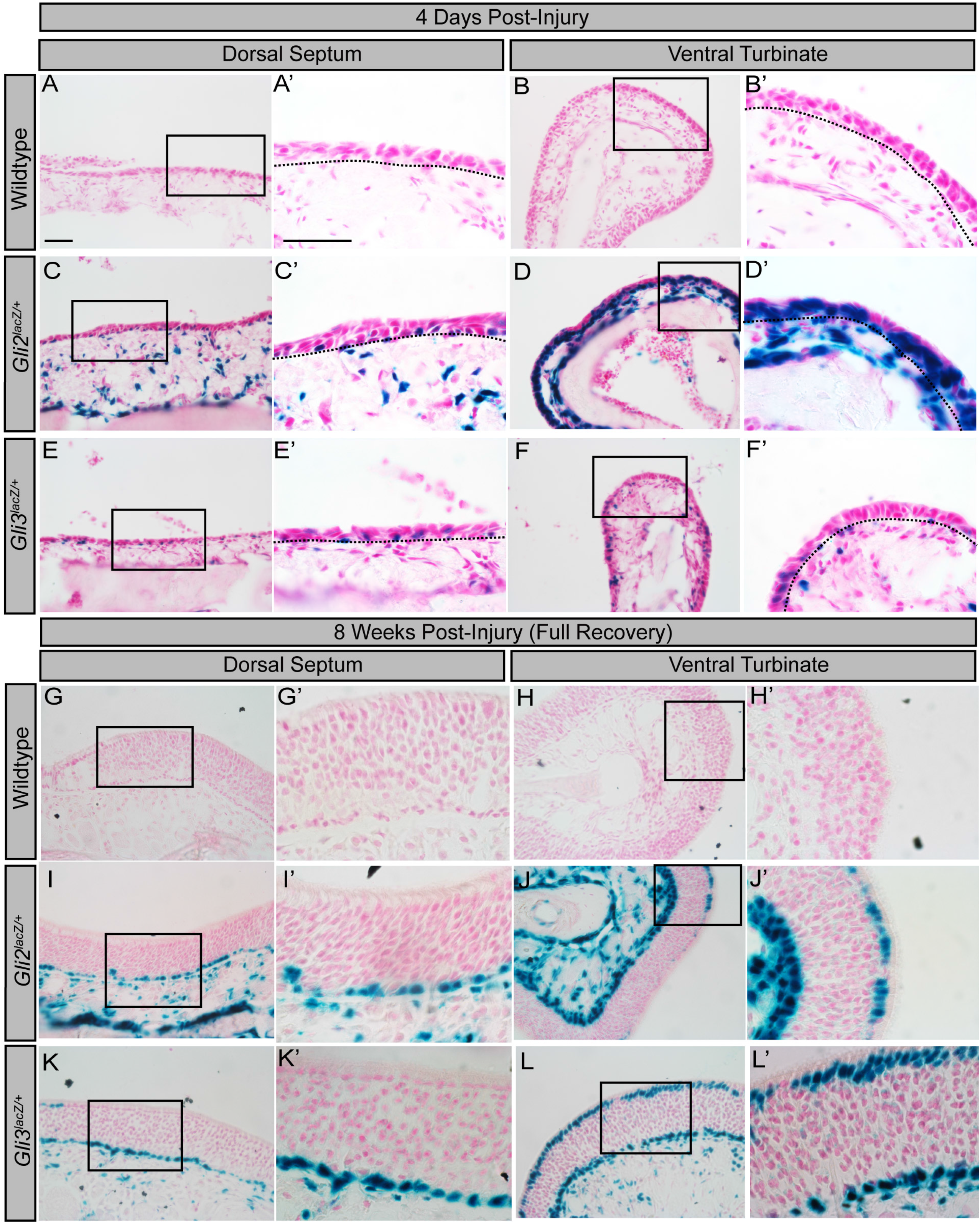
*Gli2* expression is up-regulated in ventral turbinate-associated OE at 4 days following methimazole injury. Adult wild type, *Gli2^lacZ/+^*, and *Gli3^lacZ/+^*mice were injured with 75mg/kg methimazole and allowed to recover for either 4 days (A-F’) or 8 weeks for full recovery (G-L’). X-GAL staining of coronal sections from adult wildtype (A-B’, G-H’), *Gli2^lacZ/+^* (C-D’, I-J’), and *Gli3^lacZ/+^* (E-F’, K-L’) reporter mice. Images are taken from both the dorsal septum (A, A’, C, C’, E, E’, G, G’, I, I’, K, K’) and a ventral turbinate (B, B’, D, D’, F, F’, H, H’, J, J’, L, L’).

**Figure S3.**
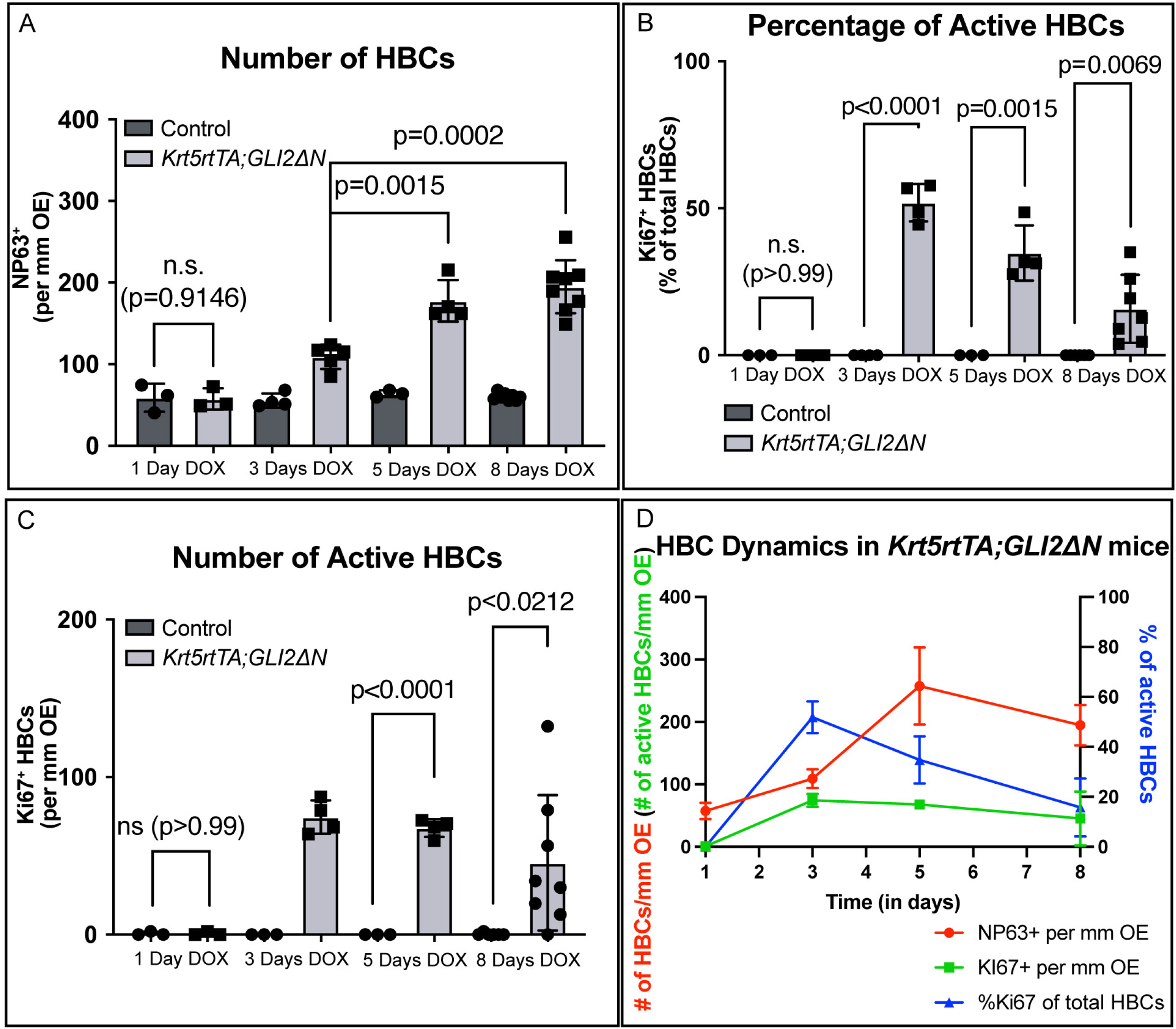
Quantitation of GLI2A-mediated HBC proliferation. (A) Quantitation of HBC number per millimeter (mm) of olfactory epithelium at 1 Day, 3 Days, 5 Days, and 8 Days following doxycycline induction. (B) Quantitation of the percentage of Ki67^+^ HBCs of total HBCs in the olfactory epithelium at 1 Day, 3 Days, 5 Days, and 8 Days following doxycycline induction. n=3 control, 3 *Krt5rtTA*; *GLI2ΔN* animals analyzed for 1 Day DOX. (C) Quantitation of Ki67^+^ HBCs per mm of olfactory epithelium at 1 Day, 3 Days, 5 Days, and 8 Days following doxycycline induction. n=4 control, n=5 *Krt5rtTA*; *GLI2ΔN* animals analyzed for 3 Days DOX. n=3 control, 4 *Krt5rtTA*; *GLI2ΔN* animals analyzed for 5 Days DOX. n=6 control, 7 *Krt5rtTA*; *GLI2ΔN* animals analyzed for 8 Days DOX. Control refers to mice containing either the *Krt5rtTA* transgene or *GLI2ΔN* transgene. Data are represented as the mean ± standard deviation. P-values were determined by a two-tailed unpaired t-test; n.s.= not significant. (D) Summary graph of HBC dynamics in *Krt5rtTA*; *GLI2ΔN* animals, red indicates the number of HBCs per mm OE, green indicates the number of Ki67^+^ HBCs per mm OE, and blue indicates the percentage of Ki67^+^ HBCs from total HBCs in the OE.

**Figure S4.**
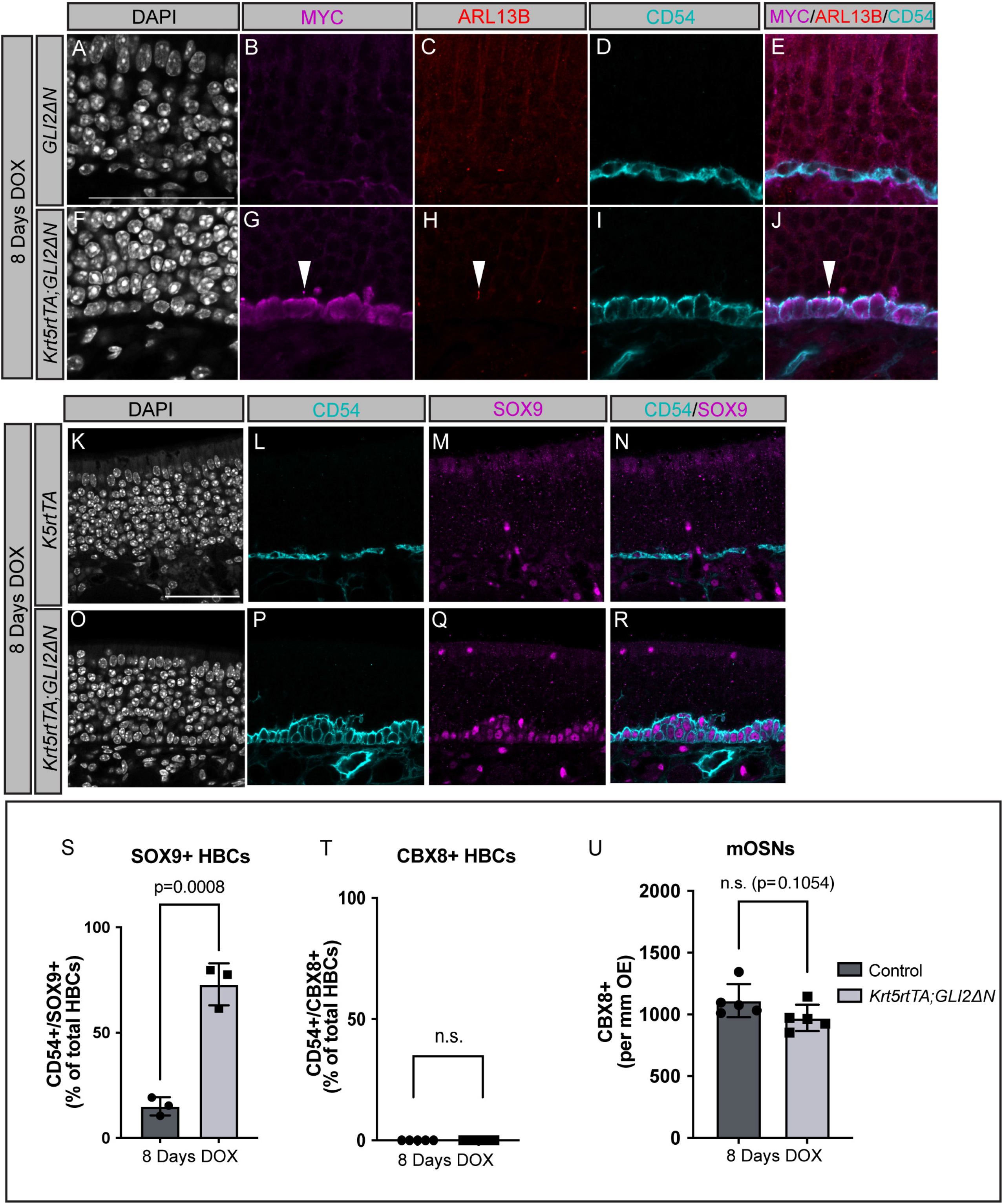
Constitutively Active GLI2 localizes to primary cilia in HBCs and increases SOX9 expression. (A-U) Mice containing the *Krt5rtTA* and *GLI2ΔN* transgenes were treated with doxycycline for 8 days. Coronal sections from adult mice were treated with antibodies directed against MYC (magenta; B, G; white arrowhead) which detects GLI2ΔN, Arl13b (red; C, H; white arrowhead) which labels primary cilia, and CD54 (cyan; D, I) which denotes HBCs. Merged MYC, Arl13b, CD54 images (E, J). DAPI denotes nuclei (A, F), scale bar (A) 50μm. (K-R) Coronal sections from adult mice treated with DOX for 8 days were analyzed with immunostaining using different OE markers. Antibodies directed against CD54 (cyan; L, P) denote HBCs, while antibody detection of SOX9 (magenta; M, Q) identifies Bowman’s gland cells. DAPI indicates nuclei (gray; K, O). Scale bar (A), 50μm. Merged CD54 and SOX9 images (N, R). Quantitation of the percentage SOX9^+^ HBCs, n=3 control and n=3 *Krt5rtTA*; *GLI2ΔN* animals analyzed (S). Quantitation of the percentage of CBX8^+^ HBCs, n=5 control and n=5 *Krt5rtTA*; *GLI2ΔN* animals analyzed (T). Quantitation of mOSNs, n=5 control and n=5 *Krt5rtTA*; *GLI2ΔN* animals analyzed (U). Control refers to mice containing either the *Krt5rtTA* or *GLI2ΔN* transgene. Data are mean ± standard deviation. P-values were determined by a two-tailed unpaired t-test; n.s.= not significant (p>0.05).

**Figure S5.**
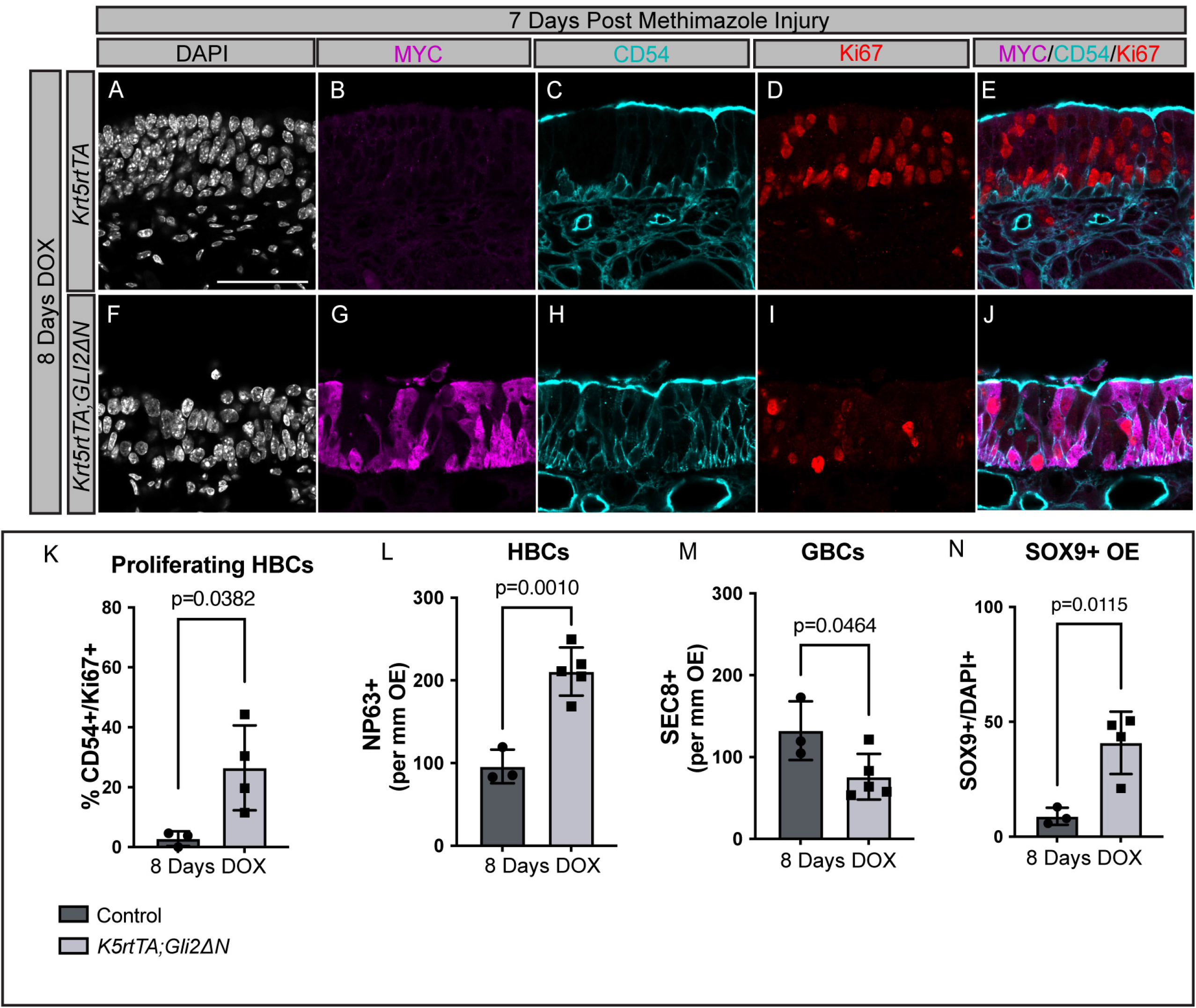
Constitutively Active GLI2 drives HBC proliferation following injury. (A-J) Coronal sections from adult mice treated with DOX for 8 days, 7 days following methimazole injury, were analyzed with immunostaining using different OE markers. Antibodies directed against MYC (magenta; B, G) denote MYC tagged GLI2ΔN, while antibodies directed against CD54 (cyan; C, H) stain HBCs. Actively cycling cells are stained with Ki67 (red; D, I), DAPI denotes nuclei (gray; A, F), MYC, CD54, Ki67 merged images (E, J). Scale bar (A), 50μm. Quantitation of the percentage of Ki67^+^ HBCs (K). Quantitation of the number of NP63+ HBCs (L), SEC8+ GBCs (M), and SOX9+ cells (N) per mm of OE. n=3 control and n=5 *Krt5rtTA*; *GLI2ΔN* animals analyzed. Control refers to mice containing either the *Krt5rtTA* or *GLI2ΔN* transgene. Data are mean ± standard deviation. P-values were determined by a two-tailed unpaired t-test; n.s.= not significant (p>0.05).

**Figure S6.**
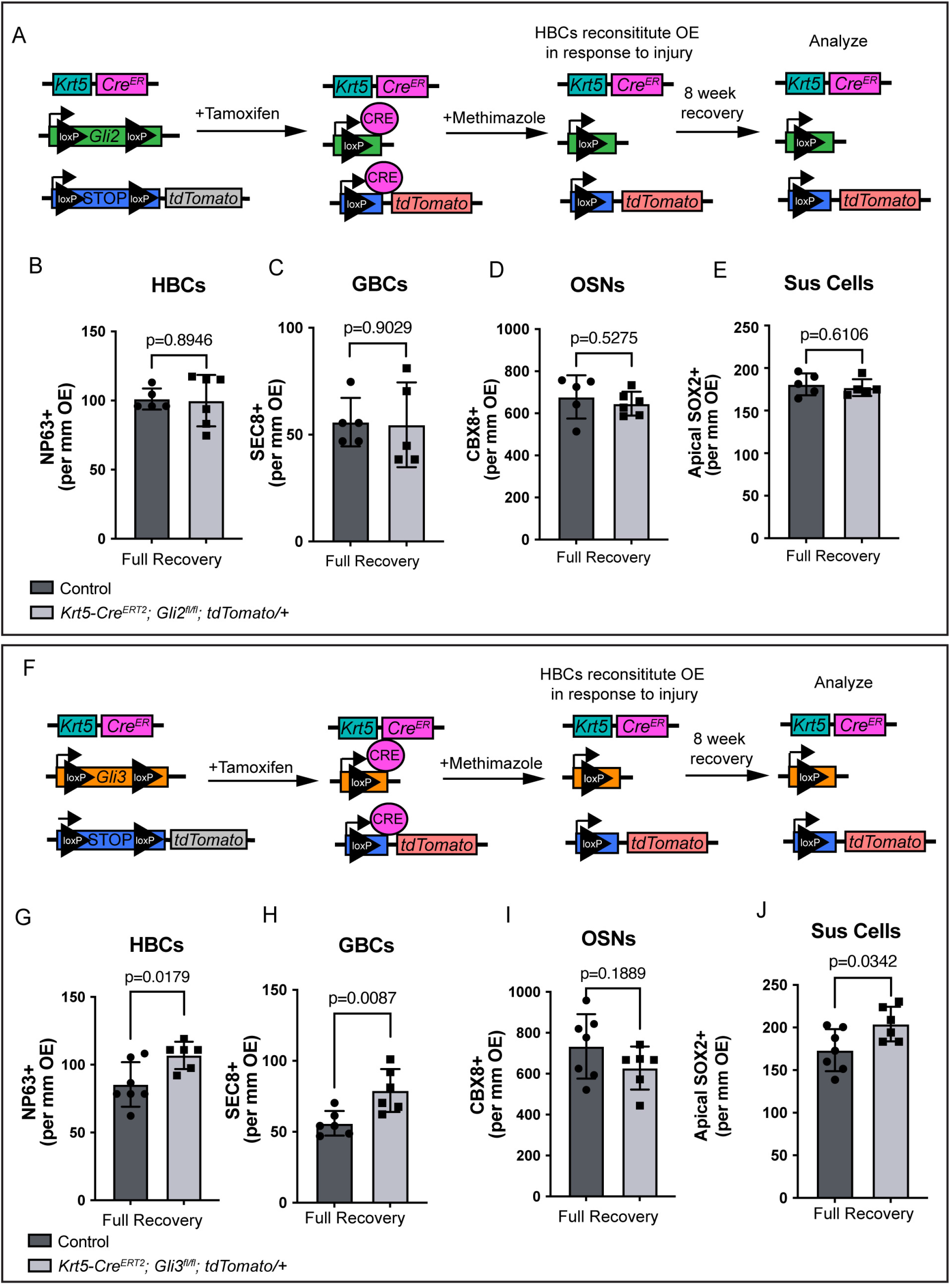
HBC-specific individual *Gli3*, but not *Gli2* deletion results in improper OE regeneration. (A, F) Cartoon of HBC-specific individual deletion of either *Gli2* or *Gli3*. Mice containing a *Krt5-Cre^ER^* Cre allele were crossed with mice carrying either *Gli2^fl/fl^* (A) or *Gli3^fl/fl^* (F), alleles in addition to a *tdTomato* reporter allele. Upon tamoxifen administration (I.P., 100mg/kg for 5 consecutive days), mice were rested for 72 hours, then injured with methimazole (I.P., 75mg/kg) and allowed to recover for 8 weeks. Quantitation of HBCs (B), GBCs (C), OSNs (D), and Sus Cells (E) per mm of OE. n=5 control and n=6 *Krt5-Cre^ER^*; *tdTomato/+*; *Gli2^fl/fl^* animals analyzed. Control refers to *Krt5-Cre^ER^*; *tdTomato/+*; *Gli2^fl/+^* mice or mice lacking the *Krt5-Cre^ER^* allele. Quantitation of HBCs (G), GBCs (H), OSNs (I), and Sus Cells (J) per mm of OE. n=7 control and n=6 *Krt5- Cre^ER^*; *tdTomato/+*; *Gli3^fl/fl^* animals analyzed. Control refers to *Krt5-Cre^ER^*; *tdTomato/+*; *Gli3^fl/+^* mice or mice lacking the *Krt5-Cre^ER^* allele. Data are mean ± standard deviation. P-values were determined by a two-tailed unpaired t-test; n.s.= not significant (p>0.05).

**Figure S7.**
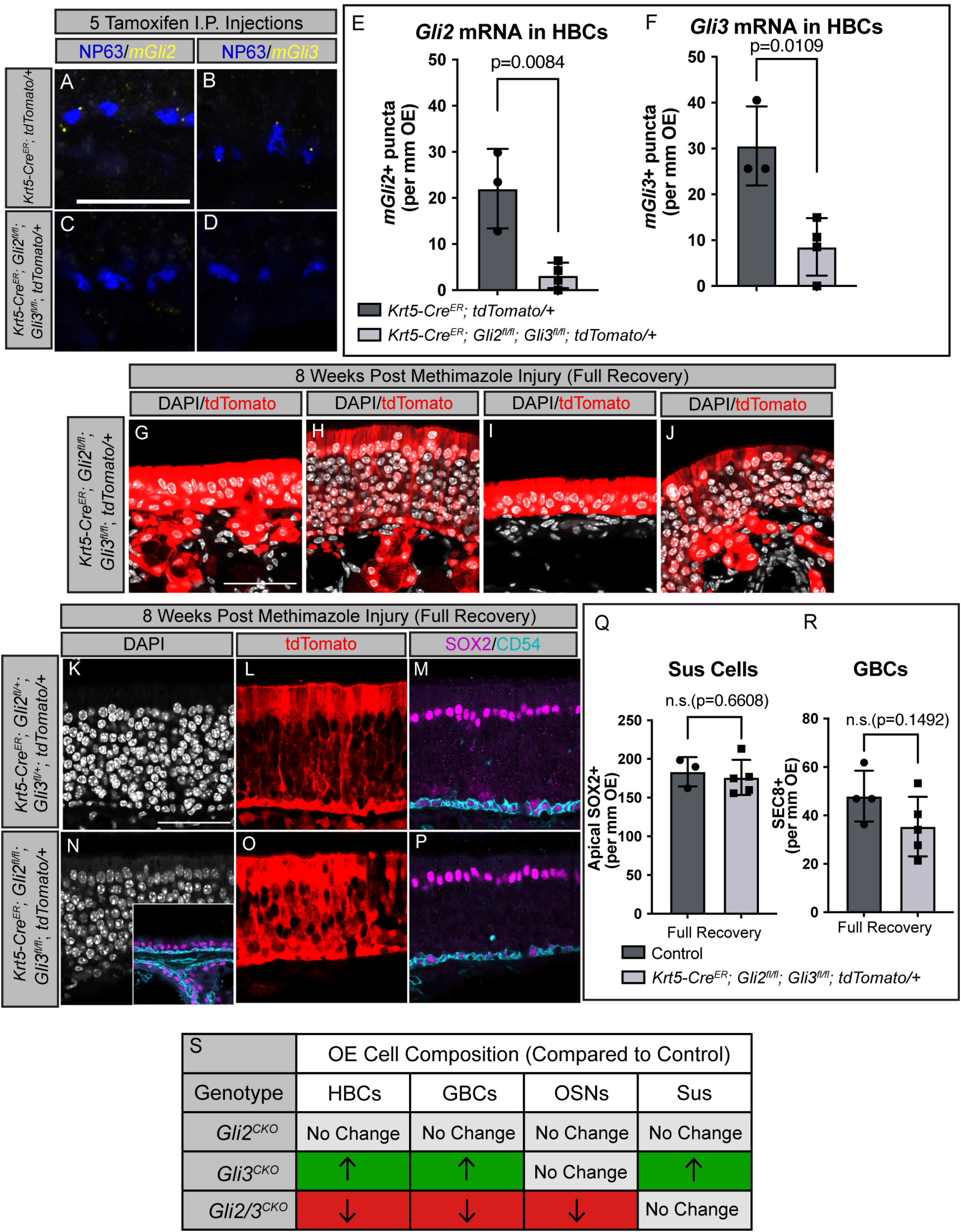
Simultaneous HBC-specific *Gli2* and *Gli3* deletion results in defective OE regeneration. (A-E) Coronal sections from adult *Krt5-Cre^ER^*; *tdTomato/+*; *Gli2^fl/fl^*; *Gli3^fl/fl^* mice collected after 5 days of consecutive tamoxifen injections (I.P., 100mg/kg). Sections were treated with mRNA probes directed against endogenous *Gli2* (yellow; A, C) and *Gli3* (yellow; B, D). Antibodies directed against NP63 demarcate HBCs (blue; A-D). Quantitation of *Gli2* (E) puncta and *Gli3* (F) puncta adjacent to NP63+ nuclei. Scale bar (A), 25μm. Data are mean ± standard deviation. P-values were determined by a two-tailed unpaired t-test; n.s.= not significant (p>0.05). (G-P) Coronal sections from adult *Krt5-Cre^ER^*; *tdTomato/+*; *Gli2^fl/fl^*; *Gli3^fl/fl^* mice collected 8 weeks following injury were analyzed with immunostaining for various OE markers. Mice lacking both *Gli2* and *Gli3* display a range of morphological phenotypes (G-J). Antibodies directed against SOX2 (magenta; M, P, inset in N) denote Sus cells apically, whereas antibodies directed against CD54 (cyan, M, P, inset in N) demarcate HBCs. DAPI denotes nuclei (gray; G-J, K, N), tdTomato marks HBCs and their progeny following injury (red; G-J, L, O), scale bar (G, K), 50μm. Quantitation of Sus cells (Q) and GBCs (R) per mm of OE. n=4 control and n=5 *Krt5-Cre^ER^*; *tdTomato/+*; *Gli2^fl/fl^*; *Gli3^fl/fl^* animals analyzed. Control refers to *Krt5-Cre^ER^*; *tdTomato/+*; *Gli2^fl/+^*; *Gli3^fl/+^* mice or mice lacking the *Krt5-Cre^ER^* allele. Data are mean ± standard deviation. P-values were determined by a two-tailed unpaired t-test; n.s.= not significant (p>0.05). (S) Table summarizing OE cell composition in *Gli2^CKO^*, *Gli3^CKO^*, and *Gli2/3^CKO^* mice compared to control mice. Green indicates increase in cell number, red indicates decrease, light gray indicates no change.

## References

Aziz, M., Goyal, H., Haghbin, H., Lee-Smith, W.M., Gajendran, M., and Perisetti, A. (2021). The Association of “Loss of Smell” to COVID-19: A Systematic Review and Meta-Analysis. Am J Med Sci 361, 216–225. 10.1016/j.amjms.2020.09.017.

Bai, C.B., Auerbach, W., Lee, J.S., Stephen, D., and Joyner, A.L. (2002). Gli2, but not Gli1, is required for initial Shh signaling and ectopic activation of the Shh pathway. Development 129, 4753–4761.

Bai, C.B., and Joyner, A.L. (2001). Gli1 can rescue the in vivo function of Gli2. Development 128, 5161–5172.

Blaess, S., Stephen, D., and Joyner, A.L. (2008). Gli3 coordinates three-dimensional patterning and growth of the tectum and cerebellum by integrating Shh and Fgf8 signaling. Development 135, 2093–2103. 10.1242/dev.015990.

Briscoe, J., and Therond, P.P. (2013). The mechanisms of Hedgehog signalling and its roles in development and disease. Nat Rev Mol Cell Biol 14, 416–429. 10.1038/nrm3598.

Chang, C.F., Chang, Y.T., Millington, G., and Brugmann, S.A. (2016). Craniofacial Ciliopathies Reveal Specific Requirements for GLI Proteins during Development of the Facial Midline. PLoS Genet 12, e1006351. 10.1371/journal.pgen.1006351.

Chen, M., Reed, R.R., and Lane, A.P. (2019). Chronic Inflammation Directs an Olfactory Stem Cell Functional Switch from Neuroregeneration to Immune Defense. Cell Stem Cell 25, 501–513 e505. 10.1016/j.stem.2019.08.011.

Corrales, J.D., Blaess, S., Mahoney, E.M., and Joyner, A.L. (2006). The level of sonic hedgehog signaling regulates the complexity of cerebellar foliation. Development 133, 1811–1821. 10.1242/dev.02351.

Diamond, I., Owolabi, T., Marco, M., Lam, C., and Glick, A. (2000). Conditional gene expression in the epidermis of transgenic mice using the tetracycline-regulated transactivators tTA and rTA linked to the keratin 5 promoter. J Invest Dermatol 115, 788–794. 10.1046/j.1523-1747.2000.00144.x.

Durante, M.A., Kurtenbach, S., Sargi, Z.B., Harbour, J.W., Choi, R., Kurtenbach, S., Goss, G.M., Matsunami, H., and Goldstein, B.J. (2020). Single-cell analysis of olfactory neurogenesis and differentiation in adult humans. Nat Neurosci 23, 323–326. 10.1038/s41593-020-0587-9.

Ermilov, A.N., Kumari, A., Li, L., Joiner, A.M., Grachtchouk, M.A., Allen, B.L., Dlugosz, A.A., and Mistretta, C.M. (2016). Maintenance of Taste Organs Is Strictly Dependent on Epithelial Hedgehog/GLI Signaling. PLoS Genet 12, e1006442. 10.1371/journal.pgen.1006442.

Fletcher, R.B., Prasol, M.S., Estrada, J., Baudhuin, A., Vranizan, K., Choi, Y.G., and Ngai, J. (2011). p63 regulates olfactory stem cell self-renewal and differentiation. Neuron 72, 748–759. 10.1016/j.neuron.2011.09.009.

Garcia, A.D., Petrova, R., Eng, L., and Joyner, A.L. (2010). Sonic hedgehog regulates discrete populations of astrocytes in the adult mouse forebrain. J Neurosci 30, 13597–13608. 10.1523/JNEUROSCI.0830-10.2010.

Genter, M.B., Deamer, N.J., Blake, B.L., Wesley, D.S., and Levi, P.E. (1995). Olfactory toxicity of methimazole: dose-response and structure-activity studies and characterization of flavin-containing monooxygenase activity in the Long-Evans rat olfactory mucosa. Toxicol Pathol 23, 477–486. 10.1177/019262339502300404.

Gong, Q., Chen, H., and Farbman, A.I. (2009). Olfactory sensory axon growth and branching is influenced by sonic hedgehog. Dev Dyn 238, 1768–1776. 10.1002/dvdy.22005.

Grachtchouk, M., Mo, R., Yu, S., Zhang, X., Sasaki, H., Hui, C.C., and Dlugosz, A.A. (2000). Basal cell carcinomas in mice overexpressing Gli2 in skin. Nat Genet 24, 216–217. 10.1038/73417.

Grachtchouk, M., Pero, J., Yang, S.H., Ermilov, A.N., Michael, L.E., Wang, A., Wilbert, D., Patel, R.M., Ferris, J., Diener, J., et al. (2011). Basal cell carcinomas in mice arise from hair follicle stem cells and multiple epithelial progenitor populations. J Clin Invest 121, 1768–1781. 10.1172/JCI46307.

Graziadei, P.P., and Graziadei, G.A. (1979). Neurogenesis and neuron regeneration in the olfactory system of mammals. I. Morphological aspects of differentiation and structural organization of the olfactory sensory neurons. J Neurocytol 8, 1–18. 10.1007/BF01206454.

Guo, Z., Packard, A., Krolewski, R.C., Harris, M.T., Manglapus, G.L., and Schwob, J.E. (2010). Expression of pax6 and sox2 in adult olfactory epithelium. J Comp Neurol 518, 4395–4418. 10.1002/cne.22463.

Herrick, D.B., Guo, Z., Jang, W., Schnittke, N., and Schwob, J.E. (2018). Canonical Notch Signaling Directs the Fate of Differentiating Neurocompetent Progenitors in the Mammalian Olfactory Epithelium. J Neurosci 38, 5022–5037. 10.1523/JNEUROSCI.0484-17.2018.

Herrick, D.B., Lin, B., Peterson, J., Schnittke, N., and Schwob, J.E. (2017). Notch1 maintains dormancy of olfactory horizontal basal cells, a reserve neural stem cell. Proc Natl Acad Sci U S A 114, E5589–E5598. 10.1073/pnas.1701333114.

Hoffman, H.J., Rawal, S., Li, C.M., and Duffy, V.B. (2016). New chemosensory component in the U.S. National Health and Nutrition Examination Survey (NHANES): first-year results for measured olfactory dysfunction. Rev Endocr Metab Disord 17, 221–240. 10.1007/s11154-016-9364-1.

Huangfu, D., Liu, A., Rakeman, A.S., Murcia, N.S., Niswander, L., and Anderson, K.V. (2003). Hedgehog signalling in the mouse requires intraflagellar transport proteins. Nature 426, 83–87. 10.1038/nature02061.

Imamura, F., and Hasegawa-Ishii, S. (2016). Environmental Toxicants-Induced Immune Responses in the Olfactory Mucosa. Front Immunol 7, 475. 10.3389/fimmu.2016.00475.

Indra, A.K., Warot, X., Brocard, J., Bornert, J.M., Xiao, J.H., Chambon, P., and Metzger, D. (1999). Temporally-controlled site-specific mutagenesis in the basal layer of the epidermis: comparison of the recombinase activity of the tamoxifen-inducible Cre-ER(T) and Cre-ER(T2) recombinases. Nucleic Acids Res 27, 4324–4327. 10.1093/nar/27.22.4324.

Joiner, A.M., Green, W.W., McIntyre, J.C., Allen, B.L., Schwob, J.E., and Martens, J.R. (2015). Primary Cilia on Horizontal Basal Cells Regulate Regeneration of the Olfactory Epithelium. J Neurosci 35, 13761–13772. 10.1523/JNEUROSCI.1708-15.2015.

Kong, J.H., Yang, L., Dessaud, E., Chuang, K., Moore, D.M., Rohatgi, R., Briscoe, J., and Novitch, B.G. (2015). Notch activity modulates the responsiveness of neural progenitors to sonic hedgehog signaling. Dev Cell 33, 373–387. 10.1016/j.devcel.2015.03.005.

Kumari, A., Ermilov, A.N., Allen, B.L., Bradley, R.M., Dlugosz, A.A., and Mistretta, C.M. (2015). Hedgehog pathway blockade with the cancer drug LDE225 disrupts taste organs and taste sensation. J Neurophysiol 113, 1034–1040. 10.1152/jn.00822.2014.

Kumari, A., Ermilov, A.N., Grachtchouk, M., Dlugosz, A.A., Allen, B.L., Bradley, R.M., and Mistretta, C.M. (2017). Recovery of taste organs and sensory function after severe loss from Hedgehog/Smoothened inhibition with cancer drug sonidegib. Proc Natl Acad Sci U S A 114, E10369–E10378. 10.1073/pnas.1712881114.

Leung, C.T., Coulombe, P.A., and Reed, R.R. (2007). Contribution of olfactory neural stem cells to tissue maintenance and regeneration. Nat Neurosci 10, 720–726. 10.1038/nn1882.

Liu, H.X., Ermilov, A., Grachtchouk, M., Li, L., Gumucio, D.L., Dlugosz, A.A., and Mistretta, C.M. (2013). Multiple Shh signaling centers participate in fungiform papilla and taste bud formation and maintenance. Dev Biol 382, 82–97. 10.1016/j.ydbio.2013.07.022.

Long, F., Zhang, X.M., Karp, S., Yang, Y., and McMahon, A.P. (2001). Genetic manipulation of hedgehog signaling in the endochondral skeleton reveals a direct role in the regulation of chondrocyte proliferation. Development 128, 5099–5108.

Madisen, L., Zwingman, T.A., Sunkin, S.M., Oh, S.W., Zariwala, H.A., Gu, H., Ng, L.L., Palmiter, R.D., Hawrylycz, M.J., Jones, A.R., et al. (2010). A robust and high-throughput Cre reporting and characterization system for the whole mouse brain. Nat Neurosci 13, 133–140. 10.1038/nn.2467.

Marczenke, M., Sunaga-Franze, D.Y., Popp, O., Althaus, I.W., Sauer, S., Mertins, P., Christ, A., Allen, B.L., and Willnow, T.E. (2021). GAS1 is required for NOTCH-dependent facilitation of SHH signaling in the ventral forebrain neuroepithelium. Development 148. 10.1242/dev.200080.

McDermott, A., Gustafsson, M., Elsam, T., Hui, C.C., Emerson, C.P., Jr., and Borycki, A.G. (2005). Gli2 and Gli3 have redundant and context-dependent function in skeletal muscle formation. Development 132, 345–357. 10.1242/dev.01537.

Memmi, E.M., Sanarico, A.G., Giacobbe, A., Peschiaroli, A., Frezza, V., Cicalese, A., Pisati, F., Tosoni, D., Zhou, H., Tonon, G., et al. (2015). p63 Sustains self-renewal of mammary cancer stem cells through regulation of Sonic Hedgehog signaling. Proc Natl Acad Sci U S A 112, 3499–3504. 10.1073/pnas.1500762112.

Mistretta, C.M., and Kumari, A. (2019). Hedgehog Signaling Regulates Taste Organs and Oral Sensation: Distinctive Roles in the Epithelium, Stroma, and Innervation. Int J Mol Sci 20. 10.3390/ijms20061341.

Morrison, E.E., and Costanzo, R.M. (1989). Scanning electron microscopic study of degeneration and regeneration in the olfactory epithelium after axotomy. J Neurocytol 18, 393–405. 10.1007/BF01190842.

Packard, A., Schnittke, N., Romano, R.A., Sinha, S., and Schwob, J.E. (2011). DeltaNp63 regulates stem cell dynamics in the mammalian olfactory epithelium. J Neurosci 31, 8748–8759. 10.1523/JNEUROSCI.0681-11.2011.

Peng, T., Frank, D.B., Kadzik, R.S., Morley, M.P., Rathi, K.S., Wang, T., Zhou, S., Cheng, L., Lu, M.M., and Morrisey, E.E. (2015). Hedgehog actively maintains adult lung quiescence and regulates repair and regeneration. Nature 526, 578–582. 10.1038/nature14984.

Petrova, R., Garcia, A.D., and Joyner, A.L. (2013). Titration of GLI3 repressor activity by sonic hedgehog signaling is critical for maintaining multiple adult neural stem cell and astrocyte functions. J Neurosci 33, 17490–17505. 10.1523/JNEUROSCI.2042-13.2013.

Petrova, R., and Joyner, A.L. (2014). Roles for Hedgehog signaling in adult organ homeostasis and repair. Development 141, 3445–3457. 10.1242/dev.083691.

Rodriguez, S., Sickles, H.M., Deleonardis, C., Alcaraz, A., Gridley, T., and Lin, D.M. (2008). Notch2 is required for maintaining sustentacular cell function in the adult mouse main olfactory epithelium. Dev Biol 314, 40–58. 10.1016/j.ydbio.2007.10.056.

Sasaki, H., Hui, C., Nakafuku, M., and Kondoh, H. (1997). A binding site for Gli proteins is essential for HNF- 3beta floor plate enhancer activity in transgenics and can respond to Shh in vitro. Development 124, 1313–1322. 10.1242/dev.124.7.1313.

Sasaki, H., Nishizaki, Y., Hui, C., Nakafuku, M., and Kondoh, H. (1999). Regulation of Gli2 and Gli3 activities by an amino-terminal repression domain: implication of Gli2 and Gli3 as primary mediators of Shh signaling. Development 126, 3915–3924. 10.1242/dev.126.17.3915.

Scales, M.K., Velez-Delgado, A., Steele, N.G., Schrader, H.E., Stabnick, A.M., Yan, W., Mercado Soto, N.M., Nwosu, Z.C., Johnson, C., Zhang, Y., et al. (2022). Combinatorial Gli activity directs immune infiltration and tumor growth in pancreatic cancer. PLoS Genet 18, e1010315. 10.1371/journal.pgen.1010315.

Schwob, J.E., Jang, W., Holbrook, E.H., Lin, B., Herrick, D.B., Peterson, J.N., and Hewitt Coleman, J. (2017). Stem and progenitor cells of the mammalian olfactory epithelium: Taking poietic license. J Comp Neurol 525, 1034–1054. 10.1002/cne.24105.

Vortkamp, A., Lee, K., Lanske, B., Segre, G.V., Kronenberg, H.M., and Tabin, C.J. (1996). Regulation of rate of cartilage differentiation by Indian hedgehog and PTH-related protein. Science 273, 613–622. 10.1126/science.273.5275.613.

Wang, B., Fallon, J.F., and Beachy, P.A. (2000). Hedgehog-regulated processing of Gli3 produces an anterior/posterior repressor gradient in the developing vertebrate limb. Cell 100, 423–434. 10.1016/s0092-8674(00)80678-9.

Wang, C., Cassandras, M., and Peng, T. (2019). The Role of Hedgehog Signaling in Adult Lung Regeneration and Maintenance. J Dev Biol 7. 10.3390/jdb7030014.

Wang, C., de Mochel, N.S.R., Christenson, S.A., Cassandras, M., Moon, R., Brumwell, A.N., Byrnes, L.E., Li, A., Yokosaki, Y., Shan, P., et al. (2018). Expansion of hedgehog disrupts mesenchymal identity and induces emphysema phenotype. J Clin Invest 128, 4343–4358. 10.1172/JCI99435.

